# Comparing the evolvability of an ancestrally reconstructed and modern adenylate kinase

**DOI:** 10.64898/2025.12.12.693990

**Authors:** MacKenzie Patterson, Hannes Ludewig, Christopher Wilson, Daniel R. Woldring, Chansik Kim, Aedan Using, Joseph Irvin, Douglas L. Theobald, Dorothee Kern

## Abstract

Directed evolution transformed protein engineering by providing a customizable framework for generating enzymes with improved catalytic performance across diverse functions. Yet modern enzymes often stall during directed evolution because populations become trapped on local fitness peaks. Researchers have suggested that ancestral enzymes offer better starting points because they are typically more thermostable. Here we propose and experimentally test an alternative explanation that does not depend on ancestral thermostability. We posit that ancestrally reconstructed sequences are unusually evolvable because they are inferred from the evolutionary lineages that survived to produce extant proteins. Less evolvable ancestors, and the trajectories emanating from them, disappeared by extinction and therefore do not contribute to reconstructed ancestors. Using thermophilic ancestral and modern adenylate kinases, we performed independent single-round selection experiments for activity in vivo and in vitro. In both settings, the ancestral enzyme tolerates a larger number of mutations, yielding more viable variants with greater genetic diversity than its modern descendants. Because mutational robustness promotes evolvability, these results support an intrinsic evolvability of reconstructed ancestral sequences that makes them superior launch points for directed evolution.

## Introduction

Ancestral sequence reconstruction (ASR) has emerged as a fundamental approach in understanding the molecular principles of protein evolution^1^. This technique has been essential in illuminating central evolutionary questions spanning proton pumps^2^, bacterial carbon fixation^3^, evolution of allostery^4^, and thermo-adaptation of enzymes^5^. Besides their role in fundamental research, ancestral proteins have accelerated protein engineering efforts in biomedicine and biotechnology^6^, including the engineering of biosensors^7^, terpene cyclases^8^, and dehalogenases^9^, among others^10,11^. This impact stems from improved expression levels, often increased (thermo)stability, and substrate promiscuity, which together make resurrected ancestral proteins unusually robust starting scaffolds for enzyme engineering.^12,13^. Importantly, these features are hallmarks of highly evolvable sequences^14^, motivating the widely held hypothesis that ancestral sequences are more evolvable relative to their modern descendants^6,12,14–16^.

Ancestral reconstructions have indeed served as starting points for successful directed evolution (DE) campaigns and rational protein design endeavors^17–19^. Even so, direct experimental evidence comparing the evolvability of ancestral and extant proteins remains limited^17^. Understanding the molecular basis for enhanced evolvability is particularly important^20^, as protein engineers routinely invest considerable effort to impart mutational robustness onto less evolvable starting points through approaches such as extensive library screening^21^, neutral drift^22,23^, chaperone overexpression^24^, or computational stability enhancement^25,26^.

Given that phenotypic robustness (a capacity to tolerate mutations) is a known driver of evolvability^27^ and that thermostability enhances robustness^28^, it has been generally assumed that the enhanced thermostability often found in ancestrally resurrected proteins underlies their increased evolvability^13,16,20,21,29–31^. If thermostability is the reason for better evolvability of ancestral proteins, one could simply use modern thermophilic proteins as starting points. Here we propose an alternative explanation: Ancestrally reconstructed sequences display higher evolvability because they are back-calculated based on only modern variants that arose from evolutionary selection of the most evolvable sequences. The vast majority of sequences went extinct because of their lower robustness.

To test our hypothesis, we compared the evolvability of a thermophilic modern adenylate kinase (Adk) and its thermophilic ancestrally reconstructed counterpart^5^ by assessing the clonal and mutational diversity that emerged after a single round of mutation and selection. We indeed observed that the ancestral enzyme tolerates a much larger number of viable variants, which also display a higher degree of sequence diversity (mutations per gene) than the modern enzyme. Since both Adks are thermophilic, these results support the hypothesis that improved evolvability of ancestrally reconstructed sequences is an inherent property that stems from the evolutionary selection uncovered by ASR rather than thermostability.

## Results

### A weak link selection platform couples adenylate kinase activity to *E. coli* fitness

We reasoned that, if ancestral Adks are more evolvable and robust than their modern counterparts, an ancestral Adk will tolerate a greater depth of clonal and sequence diversity, effectively resulting in a larger number of viable variants (Fig. 1A). Since *adk* is an essential gene and its activity has been shown to correlate with organismal fitness^32,33^, we used *adk* as a “weak link”^32^ for organismal fitness of *E. coli* strains expressing primordial and modern *adk*s and used their doubling times as a proxy for Adk activity. To accomplish this, we employed a λ red recombination and complementation placeholder plasmid (pKOCOMP-*adk*)^34^ to knock out genomic Ec*adk* and rescue Adk activity in *Escherichia coli* BW25113 with thermophilic Adks: namely ancestor 1 (ANC1) and Adk from *Caldanaerobacter subterraneus* (CsAdk), respectively (first step in scheme shown in Supplementary Fig. 1). Doubling time measurements between 10 °C and 35 °C show that below 25 °C, both strains complemented with *anc1* or Cs*adk* on the placeholder plasmid grow more slowly than their parent *E. coli* strain (Fig. 1B). This slower growth rate parallels their decreased *k*_cat_ values at these temperatures when compared to EcAdk (Fig. 1B), highlighting that Adk activity determines cellular fitness between 10 °C and 20 °C. Seeking to verify that this fitness loss originated from lower Adk activity and not from the process of genetic manipulation, we repeated the doubling time measurement of an *adk* knockout *E. coli* strain harboring Ec*adk* on the plasmid. Indeed, the recorded doubling time is within error of wild type (WT) *E. coli*. Notably, at 25 °C and above, no significant difference in growth rate is observed, implying that the speed of Adk (EcAdk: *k*_cat,40 °C_: 673.4 s^-1^, CsAdk: *k*_cat,40 °C_: 80.2 s^-1^, ANC1: *k*_cat,40 °C_: 189.1 s^-1^) is no longer the bottleneck (weak link) for growth at these temperatures.

**Fig. 1:**
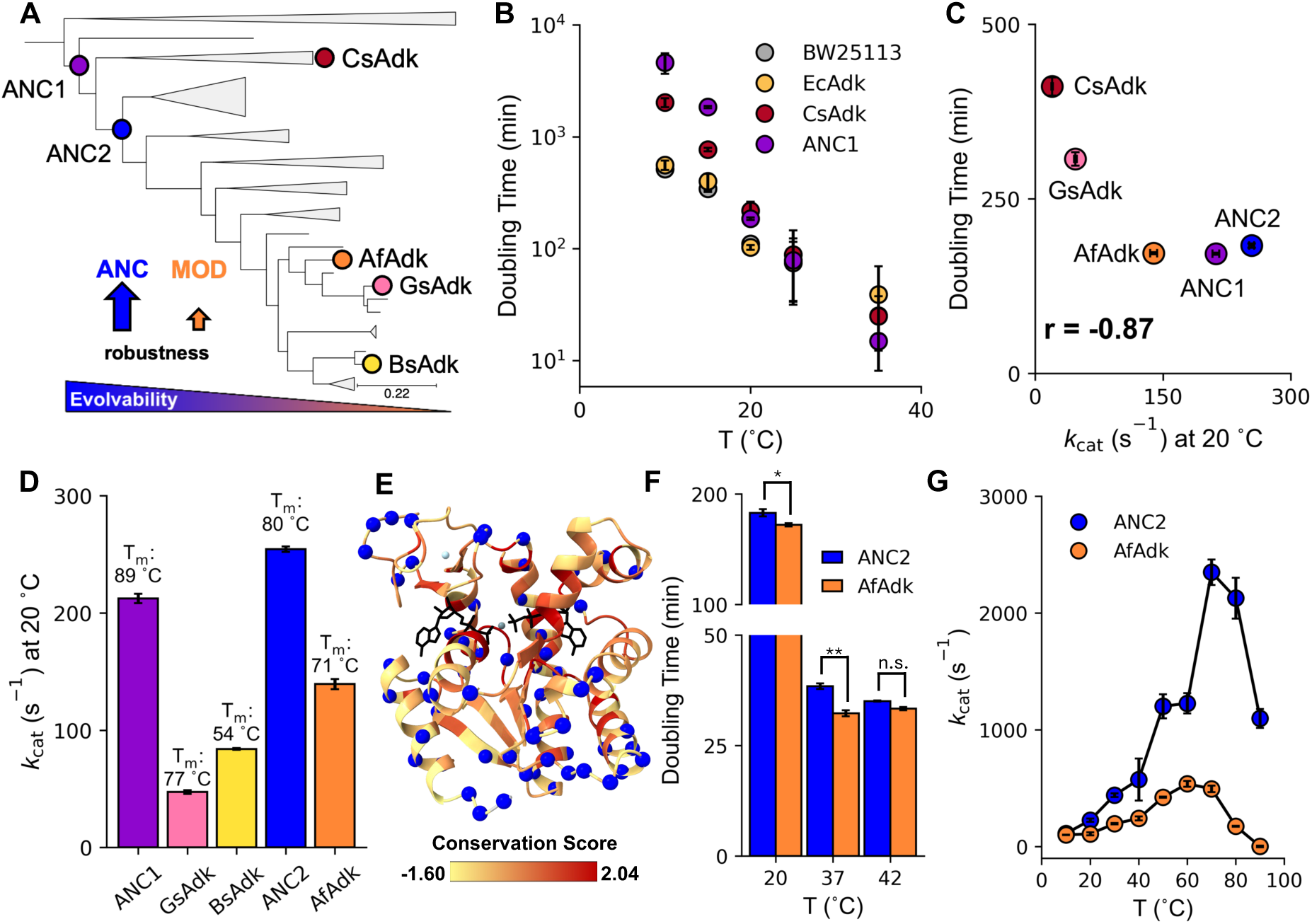
ANC2 and AfAdk are the most comparable ancestral/extant pair for *in vivo* directed evolution at low temperature. **A** Collapsed phylogenetic tree taken from Nguyen *et al.*^5^ is shown annotated with ancestral and modern Adks used in this study (Ancestor 1: ANC1, Ancestor 2: ANC2, *Caldanaerobacter subterraneus*: CsAdk, *Anoxybacillus flavithermus*: AfAdk, *Geobacillus stearothermophilus*: GsAdk, *Bacillus subtilis*: BsAdk). The hypothesized relationship between evolvability and robustness is illustrated. **B** Doubling times of *Escherichia coli* Adk (EcAdk) knockout *E. coli* BW25113 strains supplemented with indicated Adk variants on a placeholder plasmid at different temperatures. **C** Doubling times at 20 °C of knockout strains rescued with different *adks* strongly correlate with the *k*_cat_ of the rescue Adk (Pearson r = −0.87) indicating that Adk activity is rate-limiting for *E. coli* growth. **D** *k*_cat_ of Adk-catalyzed ATP production from 5 mM Mg^2+^•2ADP measured at 20 °C, via coupled assay (mean ± s.d. of n ≥ 3 replicates) and respective melting temperature T_m_ (indicated above the bar) show ANC2 and AfAdk as closest-matching ancestor-extant pair. **E** AlphaFold 3 prediction of AfAdk (cartoon) bound to Mg^2+^•ADP colored by the sequence conservation score of Adks found in the phylogenetic tree. Sequence differences between AfAdk and ANC2 are displayed as blue spheres (sequence ID 72.4%). Mg^2+^ (slate) and Zn^2+^ (light blue) are depicted as small spheres and ADP (black) as black sticks. **F** Temperature dependence of doubling times of the acceptor strain (AS; mean ± s.e. of n ≥ 3 replicates) rescued with WT ANC2 or AfAdk at different temperatures (Two-sided Kolmogorov-Smirnov test, p_20°C_ = 0.03, p_37°C_ = 0.001, p_42°C_ = 0.8; *: p < 0.05, **: p < 0.01, n.s.: not significant). **G** *k*_cat_ of ATP production from Mg^2+^•2ADP at different temperatures measured via high-pressure liquid chromatography (HPLC). All *k*_cat_ values are the average of n ≥ 3 replicate measurements ± s.e.

Emboldened by these results, we set out to produce an “acceptor strain” (AS) which could be readily transformed with high efficiency, thus being easily rescued by any sufficiently active *adk* (either a WT plasmid or a plasmid library of *adk* mutants) in a directed evolution experiment. Again, we adapted the strategy developed by Billerbeck & Panke^34^ and produced this AS using the previously created Ec*adk* knockout strain (BW25113 *adk*::Cm [pKOCOMP-Cs*adk*]) which was further transformed with the ancillary plasmid (pI-SceI). Since the placeholder plasmid we had used for genomic Ec*adk* knockout encodes Cs*adk*, the AS can survive on CsAdk activity until the strain is further transformed by the desired *adk*. The placeholder could then be removed from the resulting strain by rapid and conditional self-depletion to ensure that cell survival is solely dependent on the *adk* of interest, thereby avoiding an advantage of insufficiently active *adks*^34^. This comprehensive self-depletion is facilitated by two indispensable features of the placeholder: a heat-sensitive origin of replication, enabling plasmid curing at 42 °C, and a recognition site for the inducible restriction endonuclease, I-*Sce*I, which is encoded on an auxiliary plasmid pI-SceI (Supplementary Fig. 1 and Supplementary Table 1). To verify this method, we first transformed the AS with plasmids holding the ancestral or modern *adks* and confirmed that the doubling times of these “knock-in” strains are strongly correlated with their corresponding *k*_cat_ at 20 °C (Pearson r = −0.87; Fig. 1C).

Most importantly, this strain engineering allows selection experiments using libraries of ancestral and modern Adks. We transformed the AS with libraries of either ANC2 or one of two modern Adks, *Bacillus subtilis* (BsAdk) and *Geobacillus stearothermophilus* (GsAdk), then grew the cells at 18 °C after confirming successful removal of the placeholder plasmid. The expectation is that (i) only cells expressing active Adk variants can grow and (ii) higher-activity Adk variants will outcompete slower ones thus enrich in the culture. Sanger sequencing colonies of the selected ANC2 library indeed shows the greatest population diversity at both 42 °C and 18 °C compared to the modern Adks (Supplementary Fig. 2), indicative of the ancestral sequence being more robust. However, these first experiments have two limitations. 1) Comparisons are convoluted by differences in *k*_cat_ and thermostability (T_m_) among parental Adks: GsAdk is thermophilic but ∼8x slower at low temperatures than ANC2, while BsAdk is mesophilic and ∼3x slower than ANC2 (Fig. 1D). These discrepancies likely influence the relative response to selective pressures during the necessary 42 °C placeholder curing and at 18 °C selection for BsAdk and GsAdk library strains. 2) Sanger sequencing limits the depth (number of variants) that we could identify, thereby impeding highly quantitative comparisons of population diversity. Below we describe how we overcame these two limitations.

We endeavored to find the most similar ancestor/extant pair with respect to *k*_cat_ and T_m_ values. After extensive search, we identified Adk from *Anoxybacillus flavithermus* (AfAdk; *k*_cat_: 136 s^-1^; T_m_: 71 °C) as the best match to ANC2 (Fig. 1D). Sequence differences between the two Adks (72.4% sequence identity) are globally distributed (Fig. 1E and Supplementary Fig. 3). Notwithstanding the differences in *k*_cat_ and T_m_, the ANC2 and AfAdk WT knock-ins unveil similar fitness at 20 °C, as demonstrated by their doubling times, which are much greater (slower growth) than those exhibited at 37 °C and 42 °C where Adk activity is no longer rate-limiting for *E. coli* growth (Fig. 1F). To confirm that both enzymes are poorly adapted to 20 °C, we used a discontinuous high-pressure liquid chromatography (HPLC) assay to measure the temperature dependence of ANC2 and AfAdk *k*_cat_ values between 10 °C and 90 °C. We found they have maximal activities at 70 °C and 60 °C, respectively, with ANC2 having a much steeper increase in activity with temperature which is in agreement with other ancestral Adks^5^, but importantly show similar activities at low temperatures (Fig. 1G).

### Long-term survival selection demonstrates the enhanced evolvability of ancestral over modern adenylate kinase

Aiming to assess Adk evolvability more quantitatively, we returned to selection experiments utilizing error-prone PCR libraries of our newly identified pair, ANC2 and AfAdk. In addition to baffled culture flasks, we parallelly grew both libraries in a turbidostat^35^, which serves as a suitable, automated culturing alternative allowing for temperature regulation with continuous recording and control of optical density at 650 nm (Fig. 2A). Second, we employed a nanopore sequencing platform adapted from Karst *et al.*^36^ to quantitatively compare population composition of flasks against turbidostat cultures with much higher sequencing depth than Sanger sequencing (Supplementary Fig. 4 and Supplementary Table 1-3). We observe no differences in the survivable sequences identified from selection experiments done in flasks versus in the turbidostat (Supplementary Fig. 5). We next performed long-term selection experiments in the turbidostat, where we could continuously record doubling times, quantify bacterial population diversity via next-generation sequencing, and assess enzymatic activities of enriched individual Adks (Fig. 2A).

**Fig. 2:**
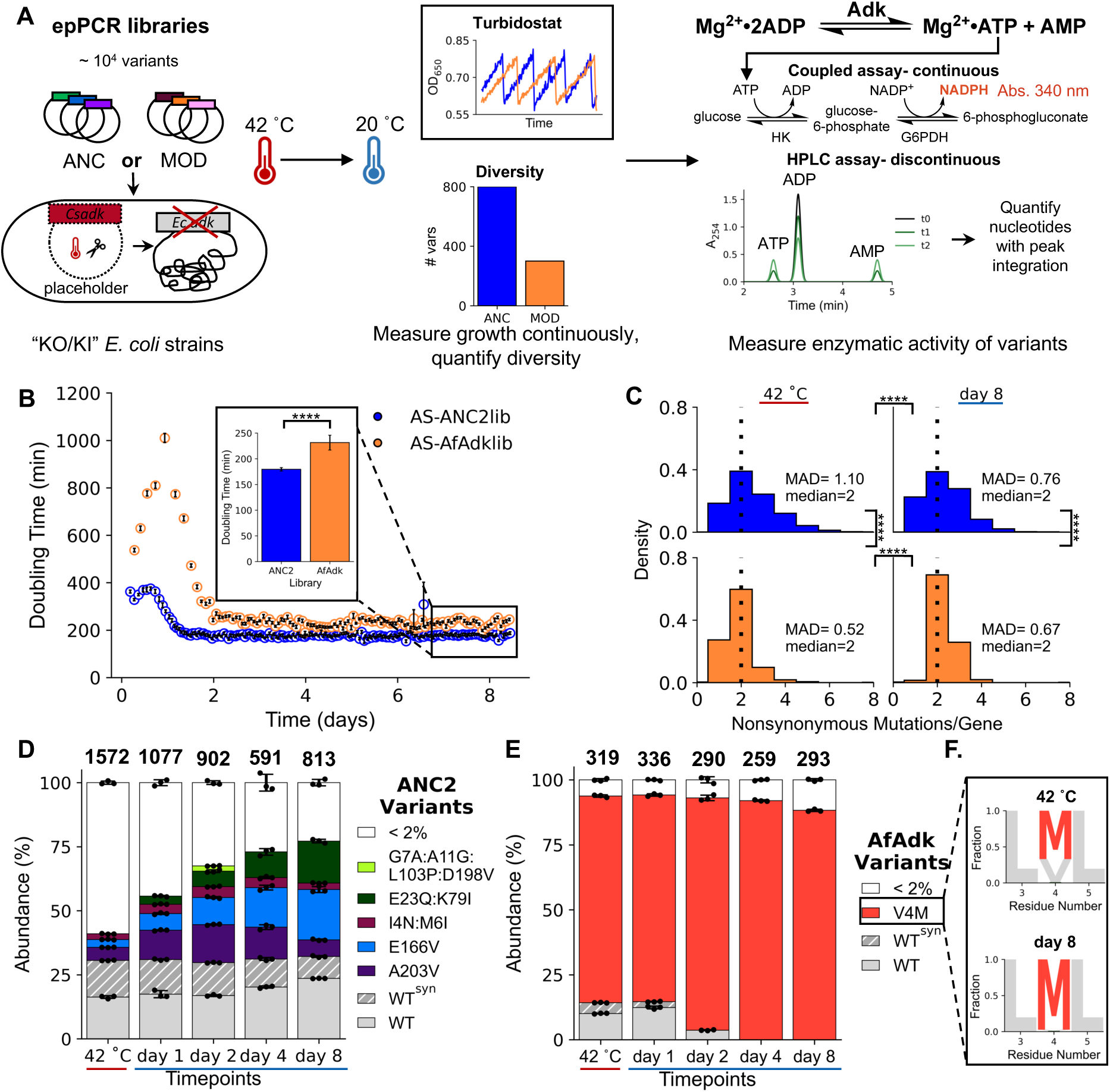
ANC2 is more robust to mutation than AfAdk at 20 °C *in vivo*. **A** Scheme of the experimental setup for *in vivo* selection of acceptor strains rescued with ancestral or modern epPCR libraries, ensuring continuous bacterial growth in log phase and continuous doubling time measurement using the Chi.Bio turbidostat^35^. Readouts for robustness are the number of viable variants, and two methods used to measure *k*_cat_ for winners are shown. **B** Doubling times of AS-ANC2lib and AS-AfAdklib strains throughout 8 days of selection at 20 °C in turbidostat, revealing a significantly faster growth of the ancestral libraries than modern libraries (Two-sided Kolmogorov-Smirnov test, p = 1.9 x 10^-31^), with an initial 2-day adaptation phase. **C** Histograms display significantly larger genetic diversity (mutations/gene) in ancestors than moderns before (at 42 °C) and after 8 days of selection at 20 °C (Two-sided Kolmogorov-Smirnov tests: p_ANC2_ = 1.04 x 10^-5^, p_AfAdk_ = 7.04 x 10^-16^, p_ANC2,AfAdk,42°C_ = 2.11 x 10^-38^, p_ANC2,AfAdk,day8_ = 5.62 x 10^-15^; ****: p < 0.001; MAD = mean absolute deviation around the median). **D-E** Throughout selection, ancestral bacterial populations are more diverse than extant ones, as measured by nanopore sequencing of AS-ANC2lib and AS-AfAdklib timepoints at 42 °C and subsequent timepoints of 20 °C selection. Numbers on top identify the total number of variants detected. **F** Sequence logo of low-abundance AS-AfAdklib variants at residues 3-5. Populations in (**D**) and (**E**) are represented as mean ± s.d. of n = 3 replicates and the average number of unique amino acid variants is indicated above each bar. Data for each replicate are provided in Supplementary Tables 2 and 3.

To ensure that selection experiment outcomes are not impacted by library diversity, we sequenced ANC2 and AfAdk libraries and quantified their diversity to be on the order of 10^4^ unique sequences. Notably, AfAdk library is slightly more diverse than the ANC2 library (Supplementary Fig. 6). After 8 days of selection AS-ANC2lib exhibited higher fitness than AS-AfAdklib as judged by doubling times of 180 and 230 min, respectively (Fig. 2B and Supplementary Fig. 7A; Kolmogorov-Smirnov p = 1.9 x 10^-31^). Characterization of population diversity implicates ANC2 as more robust, revealed by a broader distribution of the number of mutations per *adk* gene at all timepoints and temperatures (Fig. 2C). Further analysis of the nanopore sequencing reveals that not only does ANC2 tolerate more mutations per gene than AfAdk, but more importantly AS-ANC2lib continuously displays a larger number of viable Adks than AS-AfAdklib throughout our selection experiment (Fig. 2D-E and Supplementary Fig. 7B-D). The degree of enrichment of highly abundant Adks mirrors the trajectories of doubling time stabilization by each library strain. The most substantial changes in variant abundances occur during an initial ∼2-day growth adaptation, followed by a slow stabilization during the remainder of selection (Fig. 2B, D, E and Supplementary Fig. 7). Strikingly, after 8 days of selection, AS-ANC2lib contains 813 variants, 807 of which still accounted for ∼23% of the population (Fig. 2D and Supplementary Fig. 7B). In stark contrast, AS-AfAdk population is composed of 293 different variants but dominated by the single mutant AfAdk-V4M (∼80%, Fig. 2E and Supplementary Fig. 7C), despite starting from a 10^4^ variant library that was slightly larger than for ANC2. Not only is AfAdk-V4M the dominant winner of the AfAdk library selection at 20 °C, but it also serves as a compensatory mutation for almost all other low-abundance AfAdk variants in that selection (Fig. 2F). The large discrepancy in population diversities of ancestral and extant library strains is reproducible between biological replicates (Supplementary Fig. 8).

### Catalytic activities of purified variants correlate with enrichment during selection

Overall, we observe that mutations in ANC2 are distributed globally throughout the Adk structure. Contrarily, position V4 acts as a hotspot in modern Adks based on its reproducible augmentation in different modern library strains (Fig. 3A and Supplementary Fig. 2, 5, and 7). While the sequencing results convincingly exhibit greater robustness of ANC2 over AfAdk, we asked whether this trend extends to activity as well. To answer this, we used a colorimetric coupled assay to measure the Adk-catalyzed conversion from ADP to ATP and AMP of enriched ANC2 and AfAdk variants and determined that AfAdk-V4M displays a 1.3-fold increased *k*_cat_ over WT AfAdk. This small increase in activity translates into a minute 0.28 kcal/mol difference in activation free energy, which is challenging to rationalize structurally. A plausible hypothesis is a difference in hinge dynamics affecting lid opening^37^, which has been demonstrated to be the rate-limiting step^38^. These differences could be the result of alternative hinge 4 packing. In AfAdk-V4M, residue L72 (part of hinge 4) is now part of the second shell of the mutated residue 4, which resembles the arrangement in the faster ANC2 (judged by AlphaFold 3 prediction; Supplementary Fig. 9). Catalytic improvements within the AS-ANC2lib are more modest, with E166V exhibiting the greatest activity improvement of 1.1-fold over WT ANC2 (Fig. 3B, Supplementary Table 4). Despite these slight improvements, the enrichment of each highly abundant ANC2 variant is strongly correlated with its *k*_cat_ at 20 °C (Pearson r = 0.87), confirming our prediction that *E. coli* expressing faster Adks would outcompete bacteria encoding for slower Adks (Fig. 3C).

**Fig. 3:**
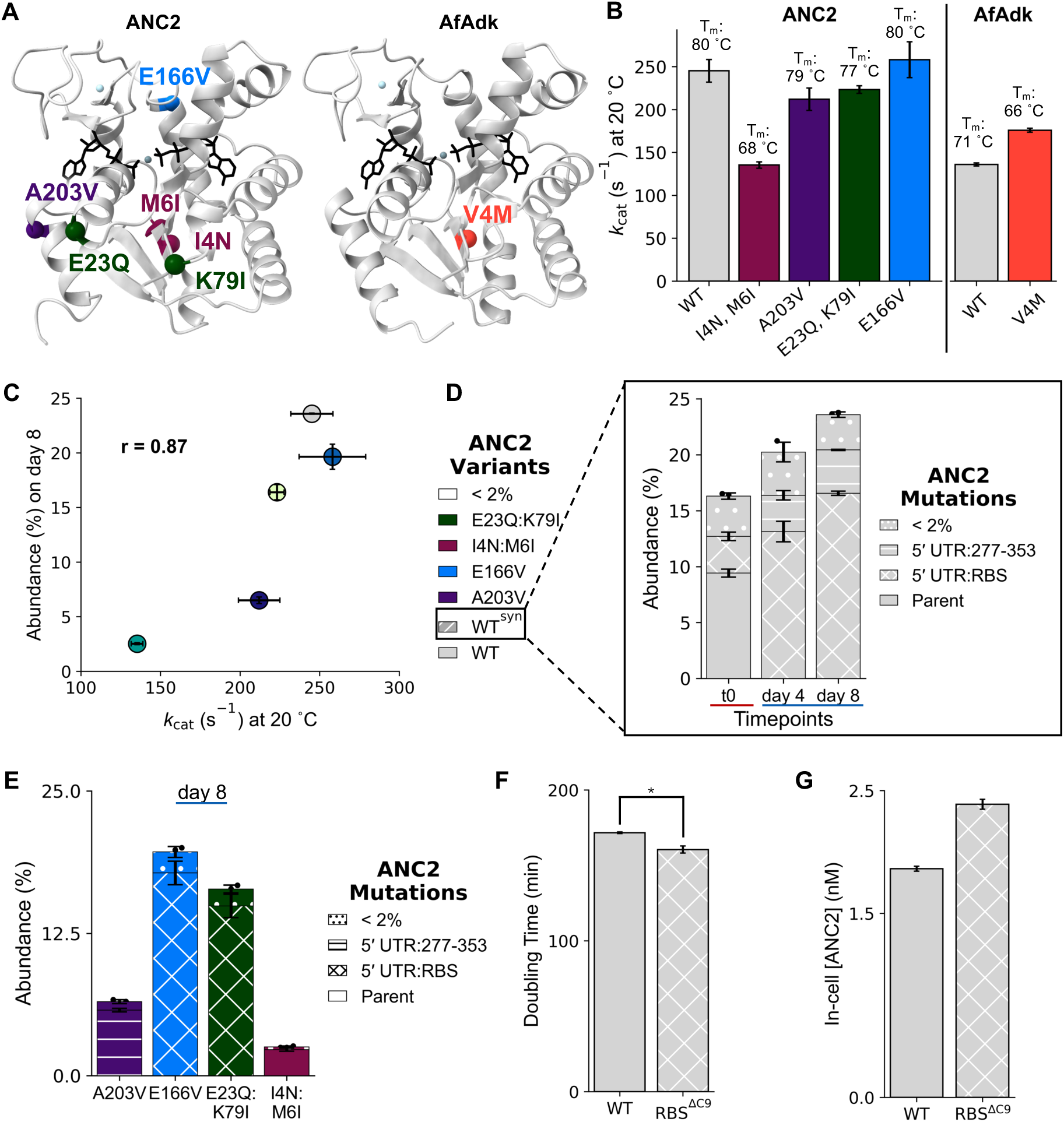
Selection *in vivo* acts on Adk activity and expression levels. **A** Mutations for each highly abundant variant after 8 days of selection are plotted onto AlphaFold 3 predictions (as colored spheres) of ANC2 or AfAdk (grey cartoon, depicted as in Fig. 1E). **B** *k*_cat_ of Adk-catalyzed ATP production from 5 mM Mg^2+^•2ADP by ANC2 and AfAdk variants enriched by selection measured via coupled assay (20 °C, mean ± s.d. of n ≥ 3 replicates, Supplementary Table 4) and melting temperature (T_m_; above bar). **C** Relative abundance of each highly abundant ANC2 variant after 8 days of selection at 20 °C plotted against its *k*_cat_ (Pearson r = 0.87) indeed shows enrichment of the fastest variants after selection. **D-E** Mutations in the 5′ UTR (5′ untranslated region) in regions 277-353 and the ribosomal binding site (RBS) are enriched in both the WT (**D**) and 3 of the 4 highly abundant ANC2 variants (**E**). Parent = wild-type vector sequence). Populations are represented as mean ± s.d. of n = 3 replicates. **F** A mutant ANC2 strain harboring the most common RBS variant, RBS^ΔC9^ (RBS numbering, see Supplementary Fig. 10H), grows faster at 20 °C than the ANC2 WT strain (Two-sided Kolmogorov-Smirnov test: p = 0.03, *: p < 0.05). **G** In-cell ANC2 concentration was determined in WT and RBS^ΔC9^ ANC2 strains (grown at 30 °C) by measuring ATP production in lysate via coupled assay at 20 °C (mean ± s.d. of n = 3; see Methods).

### Regulatory mutations reveal expression-mediated fitness effects during *in vivo* selection

Interestingly, we discovered inadvertent mutations, unpreventably introduced during epPCR library preparation. A mutation in the 5′ untranslated region (5′ UTR), selected for in clones of ANC2 WT is the most abundant and consists of a deletion of cytosine 9 in the ribosomal binding site (RBS^ΔC9^) (Fig. 3D and Supplementary Fig. 10A, B). This mutation is already present at a low level (<1%) in the starting libraries, therefore it must confer a fitness advantage during the selection. Closer inspection of the other highly abundant ANC2 sequences reveals that all but one (ANC2-I4N:M6I) contain other enriched 5′ UTR mutations (Fig. 3E and Supplementary Fig. 10C-H). Two of those enriched mutants (E166V and E23Q:K79I) contain the same RBS^ΔC9^ mutation selected for in WT (Fig. 3E and Supplementary Fig. 10E-F and H). This enrichment could possibly be caused by an elevated in-cell Adk concentration, leading to an increase in cellular Adk activity. Indeed, when we tested in-cell Adk concentration, we found that an ANC2 strain harboring a mutant RBS^ΔC9^ in its 5′ UTR displayed a higher in-cell ANC2 concentration than its WT strain and consequently effectuated a faster doubling time (Fig. 3F-G). These results agree with previous work showing that selective pressure can act on enzyme concentration rather than function to ensure survival^32,33,39^. While we note that AS-AfAdklib WT also possesses two off-target mutations in the promoter, we found that it is outcompeted by V4M, which did not contain any off-target mutations (Supplementary Fig. 11).

To further rationalize the apparent accumulation of high-abundance variants, we measured *k*_cat_ values of low-abundance sequences that were viable but were not enriched above 2% during selection. On average, these lowly populated Adks have *k*_cat_ values that are similar to or lower than WT, with one exception in ancestral (ANC2-G104R: 1.3x WT activity) and extant (AfAdk-V4M:N142D: 1.2x WT activity) libraries. Furthermore, there is no detectable relationship between their *k*_cat_ values and their abundance after 8 days of selection (Supplementary Fig. 12). Most of these variants do not contain off-target mutations. We discovered a further complication in the interpretation of the *in vivo* selection, a high degree of plasmid multimerization in biological samples (Supplementary Fig. 13). Multimerized plasmids encoded multiple origins of replication, which may allow for increased Adk expression. Alternatively, recombination of inactive Adks with active ones on the same plasmid would convolute population diversity.

### Orthogonal *in vitro* selection corroborates increased evolvability of reconstructed ancestral adenylate kinase

Despite these additional effects we found in our *in vivo* selection experiments (that are inherent to such experiments), the data robustly reveal that the ancestral enzyme is more evolvable than its modern counterpart. Nevertheless, to further buttress this key result we sought to employ an orthogonal approach and turned towards traditional *in vitro* DE. To screen Adk variant activity we used the coupled assay on lysates of *E. coli* BL21 (DE3) overexpressing epPCR variants of ANC2 or AfAdk (pET17b-library diversity: 10^4^ and Supplementary Fig. 14). To test that the coupled assay (which measures production of ATP) accurately measures ATP production by Adk, we used the addition of the Adk inhibitor AP5A as baseline and compared the ATP production of this inhibited lysate to lysate with added Mg^2+^•ADP.

The distributions of lysate Adk velocities for BL21-ANC2lib and BL21-AfAdklib (Fig. 4A) are significantly different (Kolmogorov-Smirnov p = 7.79 x 10^-6^), with BL21-ANC2lib generally exhibiting higher activities than BL21-AfAdklib (Kolmogorov-Smirnov p = 3.90 x 10^-6^). While the mean velocities of these two distributions are similar (BL21-ANC2lib_mean_ = 0.48, BL21-AfAdklib_mean_ = 0.42), their medians are strikingly different (BL21-ANC2lib_median_ = 0.34, BL21-AfAdklib_median_ = 0.13), reflective of this shift of BL21-ANC2lib toward higher activity. As a testament to higher ANC2 robustness, we not only observe fewer inactive BL21-ANC2lib Adks but importantly also observe a higher number of variants with increased activity, when compared to BL21-AfAdklib. Furthermore, BL21-AfAdklib variants exhibit a higher fraction with inactive and near-WT activities than BL21-ANC2lib.

**Fig. 4:**
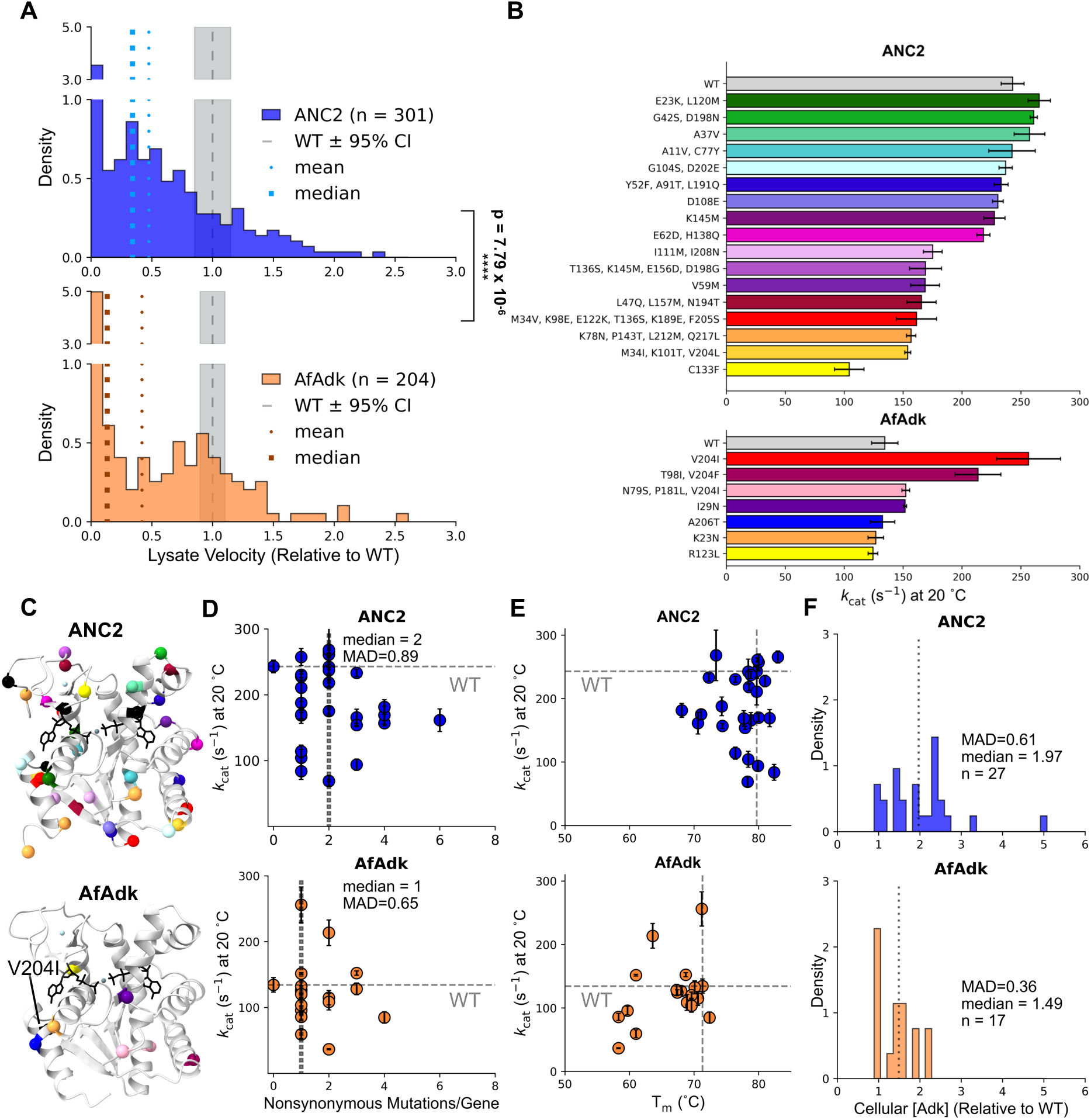
ANC2 is more robust to mutation than AfAdk in *in vitro* screening. **A** Histograms of lysate Adk activity (ATP production, coupled assay at 20 °C) show a significantly different ancestral distribution (Two-sided Kolmogorov-Smirnov test: p = 7.79 x 10^-6^), with ANC2 lysates generally being more active than AfAdk lysates (One-sided Kolmogorov-Smirnov test: p = 3.90 x 10^-6^), reflecting an increased mutational robustness in ANC2 over AfAdk (ANC2_mean_ = 0.48, ANC2_median_ = 0.34, AfAdk_mean_ = 0.42, AfAdk_median_ = 0.13). **B** Catalytic rates of ATP production by highly active Adk variants measured at 20 °C, via coupled assay (mean ± s.d. of n ≥ 2 replicates). **C** Mutations for each assayed variant are plotted using the same color as in B onto AlphaFold 3 predictions of ANC2 or AfAdk (grey cartoon, depicted as in Fig. 1E, mutations are labeled in Supplementary Fig. 15A). Mutations that appear in more than one variant are colored as black spheres. **D-E** *k*_cat_ of assayed variants at 20 °C plotted against number of mutations per gene (**D**) and T_m_ (**E**) for ANC2 and AfAdk variants. **F** Histogram showing that cellular Adk concentrations of assayed variants are significantly different (Two-sided Kolmogorov-Smirnov test: p = 0.025), with ANC2 expression levels being generally higher (One-sided Kolmogorov-Smirnov test: p = 0.013; MAD = mean absolute deviation around the median, ****: p < 0.001).

These starkly different features of ancestral and modern activity profiles mirror our *in vivo* findings and are in full agreement with our initial hypothesis that ancestral Adks would be more evolvable. We consequently expressed and purified variants with the highest lysate velocities (∼1.5-fold improved over WT) to measure *k*_cat_ values (Fig. 4B, and Supplementary Tables 5, 6). Intriguingly, we discovered another highly favored mutation site for AfAdk: residue V204, found as single mutation and in combination with other mutations (Fig. 4C and Supplementary Fig. 15A). Similarly to our *in vivo* selection experiments we found that ancestral variants harbor more mutations per gene than extant ones (Fig. 4C, D and Supplementary Fig. 15A). Overall, we observed very little perturbation in T_m_ values of ANC2 and AfAdk variants but did notice that several ANC2 variants display slightly improved thermostabilities up to 3 °C greater than WT, in contrast to only one AfAdk variant with a 1 °C increase in T_m_ (Fig. 4E). Taken together, these results suggest that ANC2 is more robust to varying amino acid changes than AfAdk.

Similarly to our *in vivo* selection, we could not find evidence that the superior robustness of ANC2 was accompanied by major catalytic improvements. There is a significant correlation between *k*_cat_ and lysate velocity for both libraries (Pearson r_ANC2_ = 0.45, p_ANC2_ = 0.02; Pearson r_AfAdk_ = 0.73, p_AfAdk_ = 5.46 x 10^-4^), confirming that lysate activity generally reports on *k*_cat_ (Supplementary Fig. 15B). The weaker correlation for ANC2 variants can be rationalized by determining Adk concentrations in screened cell lysates (using their *k*_cat_ values) and correlating these with the respective lysate velocities (Supplementary Fig. 15C). This reveals a moderate, but significant, correlation between lysate velocity and Adk concentration for ANC2 variants (Pearson r_ANC2_ = 0.55, p_ANC2_ = 0.002), but not for AfAdk variants (Pearson r_AfAdk_ = 0.39, p_AfAdk_ = 0.11). This suggests that mutations in ANC2 increase lysate activity in several cases by increasing cellular concentrations of folded Adk. This interpretation is further supported by comparing cellular Adk concentrations of ANC2 and AfAdk variants (Fig. 4F) which reveal a significant difference in their distributions (Kolmogorov-Smirnov p = 0.025), with the ancestral distribution shifted to higher cellular Adk concentration than the modern distribution (Kolmogorov-Smirnov p = 0.013). This finding is unsurprising, since our applied selective pressure acts on total cellular Adk activity, which is the product of the enzymatic rate and total active enzyme concentration in the cell. Not only does this align with our *in vivo* selection experiments but is also in agreement with studies from Shamoo and others that have demonstrated that enzyme concentration is an easily tunable parameter when adapting to selective pressures on energy metabolism^32,33,39,40^.

## Discussion

Enzymes are remarkably facile catalysts promising ever-adapting solutions to emerging fields of technology, such as synthetic biology and biocatalysis. Key to unlocking these potentials are successful DE campaigns^41^. However, DE often suffers from premature optimization plateaus^16^. Thermostability can strongly enhance the evolvability of folded proteins by increasing mutational robustness^28^. Increased stability allows proteins to tolerate neutral but destabilizing mutations without catastrophic unfolding of the scaffold, which helps evolving proteins evade local fitness maxima that can trap evolutionary trajectories^16,27,42–45^. In turn, these neutral networks enlarge the set of routes through sequence space and enable broader, more diverse exploration of the fitness landscape^27^. Given that many DE studies leverage thermophilic ancestral protein starting points for their success, it has been proposed that thermostability-driven robustness is responsible for their enhanced performance over extant scaffolds^7,9,13,16,18,20,21,29–31^. If thermostability were the primary cause, then thermostable modern enzymes, which are typically easier to obtain than ancestral reconstructions, should serve as equally strong starting points.

However, the proposed causal link between ancestral thermostability and mutational robustness still lacks conclusive experimental support. Gomez-Fernandez *et al.* raised this issue while evolving type II RuBisCOs toward higher thermostability, starting from ancestrally reconstructed and bacterial (modern) starting sequences^17^. They concluded that ancestral RuBisCO (MRPro) tolerates mutations better than the modern enzyme RubRr from *Rhodospirillum rubrum*, based on the larger fraction of active ancestral clones detected in a high-throughput lysate screen. Notably, though, the study did not comprehensively report population-level properties or the sequence diversity of the ancestral and extant starting libraries, which limits how strongly one can connect activity screening outcomes to robustness and evolvability. In contrast, Dickinson *et al.* used phage-assisted continuous evolution (PACE) to interrogate the evolution of substrate specificity in the B-cell lymphoma-2 protein family, using extant and ancestral starting points, and reported no detectable difference in sequence diversity or evolvability. It is unclear if their fragment-based Illumina sequencing approach could have resolved clonal populations well enough to detect more subtle differences^44^.

Given these conflicting results, we sought to address a focused question: does thermostability drive the elevated evolvability often observed in ancestrally reconstructed sequences? To disentangle the thermostability and evolvability of reconstructed ancestors, we initiated two independent single rounds of laboratory evolution, one *in vivo* and one *in vitro*, starting with thermophilic ancestral and modern Adks. We observed limited catalytic improvements in selected Adk variants. Due to its role as an essential gene and a limiting enzyme for bacterial growth (Fig. 1), *adk* is already optimized to maintain cellular fitness. Therefore, we did not expect to observe substantial catalytic enhancements. Instead, we chose to quantify clonal and genetic diversity after a single round of DE to assess mutational robustness as a readout for evolvability. Indeed, we consistently observed greater population size and higher sequence diversity for the ancestral Adk. These outcomes reveal an elevated mutational robustness for the ancestral enzyme that is uncoupled from ancestral thermostability. We also detected increased ancestral robustness at 42 °C (Fig. 2D-E), a temperature at which Adk activity no longer limits growth rate (Fig. 1B and F). This observation underscores that ancestral robustness can shape the outcome of laboratory evolution even when selection is not dominated by growth defects caused by Adk activity.

In addition, the ancestral variants selected *in vitro* retained, or even improved, thermostability (Fig. 4E). This property is generally beneficial for long-term evolution over additional DE rounds because it supports the accumulation of cryptic mutations. Such cryptic substitutions can potentiate epistasis, a desirable feature when tackling especially challenging protein engineering problems, because epistasis can open routes to solutions that remain inaccessible under strictly additive mutational effects^30,43,45–47^.

Our results also speak to a broader unresolved question in protein evolution: do ancestral thermostability and robustness reflect genuine pressures of ancient environments^5,48,49^, or are they a result of ASR, for example from the enrichment of consensus-like substitutions^31,50–52^?

Since our data uncouple ancestral mutational robustness from thermostability, we propose an alternative framework based on evolutionary survivorship. Modern sequences used for ASR are, by definition, descendants of past lineages that survived to the present. ASR will therefore infer ancestors that lay on evolutionary trajectories with sufficient evolvability to adapt and persist, whereas less evolvable lineages vanished by extinction and left no descendants to be sampled. Importantly, this argument does not claim that all proteins that existed billions of years ago were highly evolvable. Rather, it proposes that the specific subset of lineages now available for us to reconstruct had enough evolvability to survive. In this view, ASR preferentially recovers a “needle in the haystack” from historically explored sequence space by leveraging billions of years of evolution. Consequently, the reconstructed sequence with the highest posterior probability may not generally correspond to the most prevalent sequence at the time but instead may be one of the most evolvable sequences among those that left descendants.

We acknowledge that our exploratory attempts to explain ancestral evolvability do not yet reveal the underlying biophysical and molecular mechanisms. Mechanistic dissection would likely require substantially larger gains in catalytic rate achieved over many DE rounds, which was not the goal of this work. Even so, our study provides a practical and conceptually useful starting point for such efforts, and we hope that it encourages further testing of this framework on other reconstructed ancestral proteins.

Overall, our findings and proposed framework place ancestrally reconstructed sequences in a distinctive niche within the DE toolkit. Computational accessibility, combined with enriched evolvability that enables more thorough exploration of sequence space, makes these sequences more promising and economical DE starting points than extant sequences. Moreover, using ASR as a first-choice strategy does not preclude deploying additional DE approaches if a given engineering challenge proves remarkably stubborn.

## Methods

### Data visualization and analysis

Data visualization and statistical analysis were performed using Python (https://www.python.org). Data manipulation and analysis were performed using Pandas, NumPy, and SciPy^53–55^. Matplotlib was used for data visualization^56^. AlphaFold 3 was used to predict the ligand-bound (2ADP, Mg^2+^, Zn^2+^) structures of Adks^57^. Crystal structures and AlphaFold 3 models of Adks were visualized with ChimeraX^58^.

### Construction of error-prone PCR libraries

Error-prone PCR libraries were prepared using the GeneMorph II EZ-Clone Domain Mutagenesis kit (Agilent) following instructions provided by the manufacturer, unless stated otherwise. To achieve a high mutation rate, 1 ng of plasmid (∼ 0.15 ng of target) was amplified (35 cycles) using *adk*-flanking primers (Supplementary Table 1). DpnI digest of the EZClone reaction was performed for 2 h at 37 °C, followed by 16 h at 25 °C, and was stored at 4-10 °C until transformation (∼4-8 h). XL10-Gold Ultracompetent *E. coli* cells (Agilent) were transformed with DpnI-treated library according to the instructions provided by the manufacturer. 50 µl of the recovered culture were plated on LB agar (50 µg/mL kanamycin or 100 µg/mL ampicillin for *in vivo* and *in vitro* libraries, respectively). The remaining volume was used to inoculate a 50 mL culture of LB (respective antibiotics) and was grown overnight at 37 °C in a shaking incubator (Infors HT Multitron AJ125) at 220 rpm. Libraries were miniprepped (Qiagen) in 10X volume. DNA from 10-20 clones was Sanger sequenced (Azenta) to estimate mutation rate and confirm correct annealing to the parent plasmid. Libraries were stored at −20 °C.

### Construction of adk knockout strains

All knockout strains were made in *E. coli* (BW25113) using an adapted protocol from Billerbeck & Panke (Supplementary. S1)^34^. BW25113 *E. coli* were made chemically competent with the Hanahan method^59^ and transformed with pKOCOMP encoding *adk* from *Caldanaerobacter subterraneus* (Cs*adk*). Cell cultures were hereafter grown at 30 °C unless otherwise stated. Where applicable, antibiotics were used at the following working concetrations: ampicillin (Ap): 100 µg/ml, chloramphenicol (Cm): 34 µg/ml, gentamicin (Gm): 10 µg/ml, and kanamycin (Km): 50 µg/ml. λ red recombination^60^ was induced with 10 mM L-arabinose, and induced cells were made electrocompetent and electroporated with a chloramphenicol resistance cassette. Correct incorporation of the cassette was confirmed by colony PCR using primers flanking the knocked out genomic *Ecadk* gene, and aliquots of isolated clones from the resultant knockout strain were stored at −80 °C as 20% (v/v) glycerol stocks.

Cells from a frozen aliquot of the knockout strain were used to inoculate a culture of LB (+ Cm & Ap) and were made chemically competent^59^ for transformation with pI-SceI. The resultant strain BW25113 *adk::Cm* [pKOCOMP-Cs*adk*; pI-*SceI*] was hereafter referred to as the “acceptor strain” (AS). Isolated clones from the AS were tested for temperature and rhamnose sensitivity (for I-SceI induction) before storage as 20% (v/v) glycerol stocks at −80 °C.

A culture of 50 mL LB media was inoculated with cells from an AS glycerol stock (+ Cm, Ap & Gm), supplemented with and 1% (w/v) glucose to prevent leaky I-*Sce*I expression. The culture was grown to an OD_600_ of 0.5-0.6. Cells were harvested by centrifugation at 2,500 rcf for 10-15 min at room temperature, then washed twice in 20 mL of LB (+ Cm & Gm). The culture was resuspended in 10 mL of LB (+ Cm & Gm) and used to inoculate new 100 mL cultures of LB (+ Cm & Gm) to an OD_600_ of ∼0.05. One culture was also supplemented with ampicillin to serve as an uninduced control.

I-*Sce*I expression was induced by addition of 10 mM rhamnose to + Cm & Gm cultures when they reached an OD_600_ between 0.1 and 0.15. The induced cultures were grown until their growth started to slow relative to the uninduced control. Induced cultures were then chilled on ice for 30-60 min. All following centrifugation steps were performed at 4 °C and cells were kept on ice. All solutions for washing were pre-chilled on wet ice. Cells were pelleted by centrifugation for 10 min at 2,500 rcf and washed twice with 20 mL of water (per 50 mL of culture). Cells were then resuspended and washed in 1 mL of water, followed by 1 mL of 10% (v/v) glycerol, then resuspended in 200 µL of 10% (v/v) glycerol per electroporation (typical OD_600_ ∼30).

Electroporation was carried out in 2 mm gap cuvettes (Thermo Scientific #5520) pre-chilled on ice and with 500 ng DNA (either a homogeneous sample of WT *adk* or a heterogeneous sample of epPCR library). Cells were electroporated using a BioRad MicroPulser electroporator (EC2 setting) and immediately recovered with 800 µL of SOC media (New England Biolabs #B9020S). Qualitative pKOCOMP cleavage and negative controls were included by electroporating 2 µL of sterile water, each. Cultures were recovered in round-bottom tubes (Greiner Bio-One #187262) at 42 °C with shaking at 220 rpm for 1.5 h. After recovery, the entire volume was used to inoculate 50 mL of LB (+ Cm, Gm, & glucose) in addition to 50 µg/mL kanamycin (+ Km), except for the pKOCOMP control culture, which was grown in the presence of 100 µg/mL ampicillin (+ Cm, Gm, Ap & glucose). Cultures were grown at 42 °C with shaking at 220 rpm until reaching log phase. To check for pKOCOMP-positive clones, cultures were split to an OD_600_ of 0.2 into “+/- Ap media” (LB + Cm, Gm, Km, glucose, & +/- Ap). pKOCOMP depletion was deemed successful after a maximum of two splits. Resulting cultures were stored at −80 °C as 15% (w/v) glycerol stocks. If the pKOCOMP-depleted culture was transformed with a WT *adk* it was plated at 42 °C, and single clones were isolated and stored at −80 °C as 15% (w/v) glycerol stocks. If the culture was transformed with an epPCR library, it was immediately used to inoculate a selection experiment.

### *In vivo* selection experiments

*In vivo* selection experiments were performed in a Chi.Bio turbidostat (LABmaker, Berlin, Germany) with default settings unless otherwise stated^35^. Intrinsic turbidostat temperature control was aided by placing it in a Binder KB 115-UL incubator. All consumables (glass vials, stir bars, tubing, etc.) were purchased according to instructions provided by the manufacturer (https://chi.bio/hardware/). Before use, chamber vials and stir bars were sterilized by autoclaving, and 3-port chamber caps with O-rings were sterilized with 70% (v/v) ethanol. Filled chambers were left to equilibrate to the selection temperature before blank measurements or inoculation. The medium for selection experiments was LB (+ Cm, Gm, Km & glucose) with the addition of 25 µg/mL nystatin as an antifungal. Chambers were inoculated to an OD_600_ of ∼0.2 (OD_650_ ∼0.35-0.5). Cultures were kept in mid log phase using OD_650_ regulation dithering between OD_650_ of 0.6 and 0.8 (set point 0.7). OD_650_ was measured every minute, and intervals between dilutions were fit to a single exponential using LMFIT^61^ to calculate growth rate (*y* = *y*_0_*e*^*rt*^ where y is OD_650_, y_0_ is initial OD_650_, r is the growth rate, and t is time), then converted to a doubling time (DT) with the equation 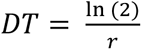 If the chamber vial was inoculated with a library, the remainder of that inoculation culture was miniprepped (Qiagen) and stored as a “42 °C”, or “pre-turbidostat” timepoint. Samples (6 mL) for genotyping were taken at least once per day and subsequently miniprepped and stored at −20 °C. For flask experiments, cultures were inoculated to an OD_600_ of 0.2 and maintained at the selection temperature in an incubator (Infors HT Multitron AJ125) with shaking at 220 rpm. Cultures were maintained in log phase (OD_600_ 0.2-0.8) manually.

### Nanopore sequencing

Sample preparation and data analysis for nanopore sequencing was adapted from Karst *et al*. using the longread_umi pipeline^36^. All primer sequences are included in Supplementary Table 1. All PCR steps were performed using Invitrogen SuperFi DNA Polymerase (Invitrogen #12351010) and Invitrogen dNTP mix (Invitrogen #18427088).

1 ng of DNA was amplified in triplicate in a 50 µL PCR reaction to obtain a linear fragment starting 86 bp upstream of the promoter and ending 79 bp downstream of the *adk* gene. Reaction components: 1X SuperFi buffer, 1 mM dNTPs, 100 nM forward primer (pSFOXB15_ont_for), 100 nM reverse primer (pSFOXB15_ont_rev), 1 ng template DNA, 0.04 units/µL SuperFi Polymerase. PCR protocol: (1) initial denaturation for 30 sec at 98 °C (2) 25 cycles of: (i) 98 °C denaturation for 10 sec, (ii) 54 °C annealing for 30 sec, (iii) 72 °C extension for 35 sec, (3) 5 min final 72 °C extension, (4) hold 10 °C. All DNA purification steps were performed with a 0.5X volume ratio of AMPure resin (Beckman Coulter #A63881). Purified fragments were eluted in 30 µL of nuclease-free water (Invitrogen #AM9939). DNA concentration was quantified using Invitrogen Quant-iT dsDNA assay kits (Thermo Scientific #Q33130 and #Q33120), according to instructions provided by the manufacturer, in opaque fluorescence 96-well plates (Corning 3993) with a SpectraMax i3x Microplate reader (Molecular Devices) or a BioTek Cytation5 Reader (Agilent). Next, each fragment was diluted to 50,000 molecules/µL (for timepoints) or 250,000 molecules/µL (for unselected libraries) and barcoded on both ends with unique molecular identifiers (UMIs) in a 50 µL PCR reaction. Reaction components: 1X SuperFi buffer, 1 mM dNTPs, 500 nM forward primer (ont_ss_dumi_for), 500 nM reverse primer (ont_ss_dumi_rev), 50,000-250,000 molecules template DNA, 0.04 units/µL SuperFi Polymerase. PCR protocol: (1) initial denaturation for 30 sec at 98 °C, (2) 2 cycles of (i) 98 °C denaturation for 10 sec, (ii) 55 °C annealing for 30 sec, (iii) 72 °C extension for 35 sec, (3) hold at 10 °C. Tagged fragments were purified and eluted in 20 µL of nuclease-free water. The entire eluate was used as the template in a 100 µL PCR reaction. Reaction components: 1X SuperFi buffer, 1 mM dNTPs, 250 nM forward primer, 250 nM reverse primer, 1 ng template DNA, 0.04 units/µL SuperFi Polymerase. PCR protocol: (1) initial denaturation for 30 sec at 98 °C, (2) 25 cycles of (i) 98 °C denaturation for 10 sec, (ii) 60 °C annealing for 30 sec, (iii) 72 °C extension for 35 sec, (3) 5 min final 72 °C extension, (4) hold 10 °C. 2 ng of the purified PCR product was re-amplified via the same PCR protocol to obtain sufficient yield for long-read sequencing.

Custom Sequencing was performed by Plasmidsaurus using Oxford Nanopore Technology to a target depth of 33X with custom analysis and annotation. The basecalled fastq files were analyzed using the longread_umi with settings longread_umi nanopore_pipeline -v 15 -s 100 -e 100 -m 750 -M 2000 - f CAAGCAGAAGACGGCATACGAGAT -F GTTATGCTATCAATCGTTGC - r AATGATACGGCGACCACCGAGATC -R GAAGGTACGCTGTATCTCAG -c 2 -p 2 - q r103_min_high_g345^36^. The output list of consensus sequences was filtered for a depth of ≥ 15X and the remaining sequences were aligned to the WT sequence using MAFFT command line wrapper ^62^ module in Biopython ^63^ with settings auto=True, inputorder=True, jtt=1, tm=2, op=1.53, ep-0.123, quiet=False. This alignment was used to extract regions from each amplicon (*e.g.,* promoter, 5′ UTR, KSD, *adk*, 5′ UTR) for easy comparison. The *adk* region of each variant was then translated using Biopython and pairwise aligned to the WT sequence for naming variants in the format w#m (w=WT ID, #=residue number, m=mutation ID). Contaminating Adk sequences were identified and removed by string comparison using RapidFuzz.^64^ Frequency of each unique Adk variant on the amino acid level was calculated by dividing the abundance of each variant by the total number of variants.

### Adk large scale expression

All *adk* overexpression constructs were designed with a C-terminal tobacco etch virus (TEV) protease cleavage/recognition site and octahistidine tag (*adk-TEV-His_8_*). Sequences were codon optimized for expression in *E. coli*, synthesized, and cloned into pET-17b by GenScript using restriction sites HindIII and KpnI. A pilot culture of LB supplemented with 100 µg/mL ampicillin was inoculated with 1 colony of BL21 (DE3) Competent *E. coli* (New England Biolabs) transformed with pET-17b-*adk-TEV-His_8_* and grown at 37 °C overnight with shaking at 220 rpm. 1-6 L of LB supplemented with 100 µg/mL ampicillin were inoculated with 20 mL/L of pilot culture and grown at 37 °C with shaking at 220 rpm until an OD_600_ of 0.6-0.8 at which point *adk-TEV-His_8_* expression was induced with 1 mM isopropyl β-D-1-thiogalactopyranoside (IPTG) for 4 h at 37 °C. Cells were harvested by centrifugation at 4,000 rpm for 15-20 min in a JLA-8.1000 rotor (Beckman Coulter) and stored at −20 °C until purification.

### Adk 96-well plate expression

BL21 (DE3) Competent *E. coli* (New England Biolabs) were transformed with 50 ng of pET-17b epPCR library according to instructions provided by the manufacturer. Alongside the libraries, two additional BL21 aliquots were transformed with 50 ng of pET-17b encoding WT *adk-TEV-His_8_* or a catalytically inactive variant, *adk-R88K-TEV-His_8_*. 10-50 µL of recovered transformation were plated on LB agar plates (100 µg/mL ampicillin) to obtain well-isolated colonies. All cell growth was done in Thermo Scientific^TM^ Nunc^TM^ Edge^TM^ 96-well, nunclon delta-treated flat-bottomed microplates (Thermo Scientific #167542) with 200 µL of LB supplemented with 100 µg/mL ampicillin. Wells were covered with CELLTREAT breathable sealing film and incubated in a MB-100 thermo-shaker. A pilot culture 96-well plate was inoculated with 1 colony per well (84 library variants, 6 WT, and 6 R88K) with controls distributed randomly across each plate. The pilot plate was grown overnight at 37 °C with shaking at 600 rpm. The next day, an expression plate was inoculated with 5 µL of pilot culture. Remaining pilot cultures were preserved as ∼10% (v/v) glycerol stocks at −80 °C. The expression plate was grown at 37 °C with shaking at 600 rpm until the average plate OD_600_ reached 0.8. All OD_600_ measurements were performed using a BioTek Cytation5 Reader (Agilent). Expression was induced in every well with 1 mM isopropyl β-D-1-thiogalactopyranoside (IPTG) for 4 h at 37 °C. OD_600_ of 5-fold diluted wells was measured to correct for cell content during data analysis. Cells were harvested by centrifugation at 1,500 rcf for 10-15 min and stored at −20 °C.

### Purification of large-scale Adk expressions

All purification steps took place at 4 °C. Cell pellets from *adk-TEV-His_8_* overexpression were resuspended in ∼20 mL of lysis buffer per gram of cell pellet. Lysis buffer was Buffer A (50 mM Tris, 300 mM NaCl, 2.5 mM β-mercaptoethanol, pH 7.5) supplemented with 1X Halt Protease inhibitor cocktail, deoxyribonuclease, and lysozyme. Resuspended cells were lysed by sonication for 10 min (20 sec on, 40 sec off, power output ≤ 40 W) and cleared by centrifugation in a JA-20 rotor (Beckman Coulter) pre-chilled to 4 °C at 18,000 rpm for 30-45 min.

The His-tagged Adk was purified via nickel-affinity chromatography on an Äkta Start. Lysate was loaded at 1-2 mL/minute on a HisTrap column (Cytiva) pre-equilibrated in Buffer A and the column was washed with Buffer A until the UV (280 nm) baselined. Subsequent wash steps were performed at 4 mL/min. To remove bound nucleotides, the column was washed with 10 column volumes (CV) of Buffer B (50 mM Tris, 1 M NaCl, 2.5 mM β-mercaptoethanol, pH 7.5). Nonspecifically bound contaminants were removed by washing with 5-10 CV of Buffer C (50 mM Tris, 300 mM NaCl, 25 mM imidazole, 2.5 mM β-mercaptoethanol, pH 7.5). His-tagged Adk was eluted with Buffer D (50 mM Tris, 300 mM NaCl, 500 mM imidazole, 2.5 mM β-mercaptoethanol, pH 7.5). His-tagged TEV protease^65^ was added to the eluate in a 1:30 TEV:total protein ratio and the sample was dialyzed overnight against 2 L of Buffer A per 30 mL of eluate in a 10 kDa MWCO Slide-A-Lyzer G2 dialysis cassette (Thermo Scientific).

The dialysate was filtered through a Whatman 0.2 µm PES membrane and loaded back onto the HisTrap column to separate cleaved Adk from the tag and TEV protease. Cleaved Adk was collected in the flow-through. Purity of Adk fractions and confirmation of successful tag cleavage by TEV protease was assessed by SDS PAGE. Adk was further purified via gel filtration on a Superdex S75 pg 16/60 or 16/600 (Cytiva) connected to an Äkta Pure and equilibrated in Buffer E (40 mM MOPS, 50 mM NaCl, 2 mM tris(2-carboxyethyl)phosphine hydrochloride, pH 7.0). Purified fractions containing Adk were combined and concentrated, if necessary, in an Amicon Ultra-15 10 kDa MWCO centrifugal filter to a concentration < 1 mM. Samples were flash frozen in liquid nitrogen and stored at −80 °C.

### Batch purification of small-scale Adk expressions

Cell pellets from 1 L *adk-TEV-His_8_* expressions were resuspended in 20 mL of Buffer A (40 mM MOPS, 300 mM NaCl, 2.5 mM β-mercaptoethanol, pH 7) supplemented with 1X Halt protease inhibitor cocktail, deoxyribonuclease, and lysozyme. Resuspended cells were lysed by sonication using a microtip for 2 min (20 sec on, 40 sec off, power output ≤ 21 W). Lysate was cleared by centrifugation in a JA-14.5 rotor (Beckman Coulter) pre-chilled to 4 °C for 1 h at 8,000 rpm. His-tagged Adk was purified in batch by cobalt affinity. Cleared lysate was filtered directly onto 5 mL of TALON affinity resin (Takara Bio) pre-equilibrated in Buffer A, then incubated for 1 h with gentle agitation at 80 rpm in a shaking incubator (Infors HT Multitron AJ125) at 15 °C or on a rotating platform at room temperature.

All centrifugation steps were performed for 5-10 min at 3,000 rcf and 10 °C. After decanting the supernatant, the resin was washed in > 10 CV of Buffer B (40 mM MOPS, 1 M NaCl, 2.5 mM β-mercaptoethanol, pH 7.0) to remove bound nucleotides. Nonspecifically bound proteins were washed with > 10 CV of Buffer C (40 mM MOPS, 300 mM NaCl, 5 mM imidazole, 2.5 mM β-mercaptoethanol, pH 7.0). The resin was washed then with > 10 CV of Buffer A to remove all imidazole, resuspended in 1 mL of Buffer A, and an on-resin tag cleavage was initiated by adding 300 µg of His-tagged TEV protease^65^. Cleavage proceeded at 4 °C overnight with gentle agitation on a rotating platform.

Cleavage and purity were assessed via SDS-PAGE. Samples were always ≥ 95% pure (data not shown). The samples were then filtered with a Whatman 0.2 µm PES membrane to remove residual resin and slowly mixed with a concentrated stock of tris(2-carboxyethyl)phosphine hydrochloride (250-500 mM) in Buffer A to preserve reducing agent in the samples. The samples were concentrated in Amicon Ultra-0.5 10 kDa MWCO centrifugal filters (Millipore Sigma) to concentrations ranging from 200-500 µM. Precipitated debris was removed by centrifuging each sample at 10 °C for 10 min at 14,000 rcf and transferring the supernatant into a new tube. Batch-purified Adks occasionally had high 260/280 values when measured by NanoDrop (Thermo Scientific NanoDrop One Microvolume UV-Vis Spectrophotometer or NanoDrop 2000c), thereby making A_280_ readings inaccurate, so a Bradford assay (Thermo Scientific #A55866) was used to quantify protein concentration for these samples. The purified samples were then assayed immediately, after which they were flash frozen in liquid nitrogen and stored at - 80 °C.

### Coupled assay of purified Adk

Enzymatic activity of Adks was measured in triplicate using a spectroscopic assay coupled to the reduction of nicotinamide adenine dinucleotide phosphate (NADP^+^ to NADPH). NADPH formation was measured at 340 nm using either a SpectraMax i3x Microplate Reader (Molecular Devices) or a BioTek Cytation5 Reader (Agilent). Assay buffer was 40 mM MOPS, 80 mM KCl, pH 7.0. Assays were performed in a 100 µL volume containing 10 units of hexokinase from yeast (Millipore Sigma), 10 units of glucose-6-phosphate dehydrogenase from *Leuconostoc mesenteroides* (Fisher Scientific), 0.5 mM NADP^+^, 5 mM glucose, 5 mM MgCl_2_, 5 mM adenosine diphosphate (ADP), 0.3 mg/mL bovine serum albumin (BSA), and 1-5 nM Adk in assay buffer. A master mix containing assay buffer, hexokinase, glucose-6-phosphate dehydrogenase, NADP^+^, and BSA was incubated in a covered microplate (Corning 3994) at the assay temperature for ≥ 15 min. After incubation, glucose and then Mg^2+^•ADP were added to each well. All reagents were also incubated at the assay temperature for ≥ 15 min. Between addition of each reagent there was an additional incubation period of 5 min, during which a baseline A_340_ measurement was recorded. The reaction was initiated with Adk and measured for 30 min. Background reactions were “initiated” with buffer instead of Adk and subtracted from + Adk data. The linear portion of each reaction was used to calculate a slope of NADPH formation vs. time using the correction coefficient 0.0032 or 0.0031 µM^-1^cm^-1^ for the SpectraMax and Cytation5 plate readers, respectively, and divided by Adk concentration to calculate a *k*_cat_.

### Coupled assay of the acceptor strain expressing Adk variants

Triplicate samples containing 5 x 10^7^ cells were pelleted by centrifugation for 5 min at 5,000 rcf (4 °C). Pellets were resuspended in 1 mL of assay buffer and sonicated for 20 sec using a microtip (10 sec on, 20 sec off, power output ≤ 15 W) then cleared by centrifugation at 4 °C for 30 min at 16,900 rcf. 40 µL of lysate per reaction was used in the assay. The coupled assay was performed similarly as to what was described for purified Adks, but with 1 mM NADP^+^, and lysate was incubated with the master mix and so the reaction was initiated by addition of Mg^2+^•ADP. 500 µM of the Adk inhibitor P^1^,P^5^- Di(adenosine-5′) pentaphosphate (Ap5A) was also included for lysates as a negative control for basal lysate activity. The linear portion of each reaction was used to calculate a slope of NADPH formation vs. time using the correction coefficient 0.0032 or 0.0031 µM^-1^cm^-1^ for the SpectraMax and Cytation5 plate readers, respectively, and background-corrected slopes were divided by the *k*_cat_ to calculate in-cell Adk concentration.

### Coupled assay for *in vitro* lysate screen

To prepare *in vitro* lysates, each sample was resuspended in 40 µL of 1X BugBuster (Millipore Sigma) diluted in assay buffer and incubated at room temperature with shaking in a MB-100 thermo-shaker at 600 rpm for 1 h. The lysate was cleared by centrifugation at 1,500 rcf (4 °C) for 10-15 min and 10 µL of 1/4,900-fold diluted lysate was used in activity assays.

The coupled assay was performed similarly as to what was described for purified Adks, but with 1 mM NADP^+^, and lysate was incubated with the master mix and so the reaction was initiated by addition of Mg^2+^•ADP. Wells expressing the catalytically dead R88K variant were used as negative controls for basal lysate activity. The linear portion of each reaction was used to calculate a slope of NADPH formation vs. time using the correction coefficient 0.0032 or 0.0031 µM^-1^cm^-1^ for the SpectraMax and Cytation5 plate readers, respectively, and background-corrected slopes were divided by the final measured OD_600_ to correct for cell content per well. All velocities were normalized to WT activity. Only wells that were induced at OD_600_ between 0.2-1 were analyzed to reduce false negatives from variants that did not express well due to being induced at an OD_600_ that was too low or too high. For wells that were genotyped, Adk concentrations were calculated by dividing the velocities by *k*_cat_.

### Discontinuous HPLC assay

High-pressure liquid chromatography (HPLC) assays to assess activity-temperature profiles were performed from 10-90 °C (10 °C increments). Adk was pre-incubated for 15 min at the target temperature before reaction initiation with 5 mM Mg^2+^•ADP. Between 10-40 °C, 10 nM and 20 nM of ANC2 and AfAdk, respectively, were used and between 50-90 °C 5 nM and 10 nM of ANC2 and AfAdk, respectively. 10 µL timepoints (20, 40, 60, 90, and 120 s) at each temperature were collected and quenched with 10% (v/v) TCA (10 µL) and cleared by centrifuging in a Spin-X centrifugal tube filter (Costar #8160) for 10 min at 5,000 rpm. The pH was neutralized by addition of 20 µL of 1.5 mM HEPES, pH 8.0. Samples were stored at −20 °C until analysis.

Nucleotide amounts in each sample were analyzed using an HPLC system (Agilent Infinity 1260) with an analytical HPLC column (ACE; EXL-129-2502U, i.d. 2.1 mm, length 250 mm, C18-AR, 5 Å pore size). Isocratic elution with a 100 mM potassium phosphate, pH 6.0 mobile phase was used for nucleotide separation. ATP, ADP, and AMP were individually integrated (ChemStation) and their ratios were used to calculate nominal concentration considering a total of 5 mM of nucleotides. The reaction velocity at each temperature was determined by linear regression (Prism 10.1.1 GraphPad Software) of ATP production over time and *k*_cat_ was calculated by dividing velocity by enzyme concentration. Depicted errors are the propagated standard error of the slope obtained from linear regression. Control experiments using only buffer or Mg^2+^•ADP were performed to examine residual contaminants.

### Determination of melting temperatures

Melting temperatures were determined using a thermal shift assay conducted in a QuantStudio 3 Real-Time PCR System (Applied Biosystems) according to manufacturer’s instructions. 5-20 µM of Adk was combined with 5X SYPRO Orange (Invitrogen #S6650) in assay buffer (20 µL total volume) in a MicroAmp Optical 96-well reaction plate (Applied Biosystems #N8010560). Melt curves were generated by measuring fluorescence intensity from 20-95 °C. The inflection point of the melting curve was used to determine the melting temperature.

## Data Availability

Raw Nanopore data is deposited on the Sequence Read Archive with accession number PRJNA1404875.

## Code Availability

The code that supports the findings of this study is available on Zenodo with doi: 10.5281/zenodo.18301099.

## Acknowledgements

We would like to thank Michael Sennett, Ricardo Pádua, Lufan Xiao and other members of the Kern and Theobald labs for valuable discussions. Additionally, we are grateful for the advice of Sven Panke and Sonja Billerbeck concerning the acceptor strain, Søren Karst regarding sample preparation for and implementation of their nanopore data analysis pipeline, and Harrison Steel concerning the Chi.Bio turbidostat.

## Author contributions

M.P., H.L., C.W., D.R.W., C.K., D.L.T., and D.K. contributed to the design and conceptualization of the project. M.P., H.L., C.W., D.R.W., and C.K. made knockout and WT knock-in cell strains. M.P., H.L., D.R.W., C.K., and J.I. measured WT acceptor strain doubling times. M.P. made error-prone PCR libraries. M.P. and H.L. performed *in vivo* experiments. M.P. prepared and analyzed all nanopore sequencing data. M.P. performed *in vitro* selection experiments. M.P., H.L., C.K., and A.U. expressed, purified, and measured enzymatic activity and thermostability WT and variant Adks. M.P., H.L., C.K., and A.U. expressed, purified, and measured enzymatic activity and thermostability WT and variant Adks. H.L. performed the HPLC assay of Adks. M.P., H.L., and D.K. wrote the manuscript. All authors reviewed the manuscript.

## References

1. Linus Pauling, Emile Zuckerkandl, Thormod Henriksen, & Rolf A. Løvstad. Chemical Paleogenetics. Molecular ‘Restoration Studies’ of Extinct Forms of Life. Acta Chemica Scandinavica https://doi.org/10.3891/acta.chem.scand.17s-0009 (1963) doi:10.3891/acta.chem.scand.17s-0009.

2. Finnigan, G. C., Hanson-Smith, V., Stevens, T. H. & Thornton, J. W. Evolution of increased complexity in a molecular machine. Nature 481, 360–364 (2012).

3. Schulz, L. et al. Evolution of increased complexity and specificity at the dawn of form I Rubiscos. Science 378, 155–160 (2022).

4. Hadzipasic, A. et al. Ancient origins of allosteric activation in a Ser-Thr kinase. Science 367, 912–917 (2020).

5. Nguyen, V. et al. Evolutionary drivers of thermoadaptation in enzyme catalysis. Science 355, 289–294 (2017).

6. Prakinee, K., Phaisan, S., Kongjaroon, S. & Chaiyen, P. Ancestral Sequence Reconstruction for Designing Biocatalysts and Investigating their Functional Mechanisms. JACS Au 4, 4571–4591 (2024).

7. Whitfield, J. H. et al. Construction of a robust and sensitive arginine biosensor through ancestral protein reconstruction. Protein Science 24, 1412–1422 (2015).

8. Schriever, K. et al. Engineering of Ancestors as a Tool to Elucidate Structure, Mechanism, and Specificity of Extant Terpene Cyclase. J. Am. Chem. Soc. 143, 3794– 3807 (2021).

9. Babkova, P., Sebestova, E., Brezovsky, J., Chaloupkova, R. & Damborsky, J. Ancestral Haloalkane Dehalogenases Show Robustness and Unique Substrate Specificity. ChemBioChem 18, 1448–1456 (2017).

10. Gutierrez-Rus, L. I. et al. Ancestral <em>versus</em> Modern Substrate Scope in Family-1 Glycosidases. bioRxiv 2024.04.11.589065 (2024) doi:10.1101/2024.04.11.589065.

11. Keser, M. et al. Stable and Promiscuous Galactose Oxidases Engineered by Directed Evolution, Atomistic Design, and Ancestral Sequence Reconstruction. ACS Synth. Biol. 14, 239–246 (2025).

12. Spence, M. A., Kaczmarski, J. A., Saunders, J. W. & Jackson, C. J. Ancestral sequence reconstruction for protein engineers. Current Opinion in Structural Biology 69, 131–141 (2021).

13. Alcalde, M. When directed evolution met ancestral enzyme resurrection. Microb. Biotechnol. 10, 22–24 (2017).

14. Tokuriki, N. & Tawfik, D. S. Protein Dynamism and Evolvability. Science 324, 203–207 (2009).

15. Tokuriki, N. & Tawfik, D. S. Stability effects of mutations and protein evolvability. Current Opinion in Structural Biology 19, 596–604 (2009).

16. Goldsmith, M. & Tawfik, D. S. Enzyme engineering: reaching the maximal catalytic efficiency peak. Current Opinion in Structural Biology 47, 140–150 (2017).

17. Gomez-Fernandez, B. J. et al. Directed -in vitro-evolution of Precambrian and extant Rubiscos. Scientific Reports 8, 5532 (2018).

18. Risso, V. A. et al. De novo active sites for resurrected Precambrian enzymes. Nature Communications 8, 16113 (2017).

19. Gumulya, Y. et al. Engineering highly functional thermostable proteins using ancestral sequence reconstruction. Nature Catalysis 1, 878–888 (2018).

20. Tokuriki, N. & Tawfik, D. S. Stability effects of mutations and protein evolvability. Curr. Opin. Struct. Biol. 19, 596–604 (2009).

21. Cole, M. F. & Gaucher, E. A. Utilizing natural diversity to evolve protein function: applications towards thermostability. Current Opinion in Chemical Biology 15, 399–406 (2011).

22. Bershtein, S., Segal, M., Bekerman, R., Tokuriki, N. & Tawfik, D. S. Robustness–epistasis link shapes the fitness landscape of a randomly drifting protein. Nature 444, 929–932 (2006).

23. Hart, K. M. et al. Thermodynamic System Drift in Protein Evolution. PLOS Biology 12, e1001994 (2014).

24. Tokuriki, N. & Tawfik, D. S. Chaperonin overexpression promotes genetic variation and enzyme evolution. Nature 459, 668–673 (2009).

25. Goldenzweig, A. et al. Automated Structure- and Sequence-Based Design of Proteins for High Bacterial Expression and Stability. Molecular Cell 63, 337–346 (2016).

26. Goldenzweig, A. & Fleishman, S. Principles of Protein Stability and Their Application in Computational Design. Annu. Rev. Biochem. 87, 1–25 (2018).

27. Wagner, A. Robustness and evolvability: a paradox resolved. Proceedings of the Royal Society B: Biological Sciences 275, 91–100 (2007).

28. Bloom, J. D., Labthavikul, S. T., Otey, C. R. & Arnold, F. H. Protein stability promotes evolvability. Proceedings of the National Academy of Sciences 103, 5869–5874 (2006).

29. Wheeler, L. C., Lim, S. A., Marqusee, S. & Harms, M. J. The thermostability and specificity of ancient proteins. Current Opinion in Structural Biology 38, 37–43 (2016).

30. Harms, M. J. & Thornton, J. W. Evolutionary biochemistry: revealing the historical and physical causes of protein properties. Nature Reviews Genetics 14, 559–571 (2013).

31. Trudeau, D. L., Kaltenbach, M. & Tawfik, D. S. On the Potential Origins of the High Stability of Reconstructed Ancestral Proteins. Molecular Biology and Evolution 33, 2633–2641 (2016).

32. Couñago, R., Chen, S. & Shamoo, Y. In Vivo Molecular Evolution Reveals Biophysical Origins of Organismal Fitness. Molecular Cell 22, 441–449 (2006).

33. Couñago, R. & Shamoo, Y. Gene replacement of adenylate kinase in the gram-positive thermophile Geobacillus stearothermophilus disrupts adenine nucleotide homeostasis and reduces cell viability. Extremophiles 9, 135–144 (2005).

34. Billerbeck, S. & Panke, S. A genetic replacement system for selection-based engineering of essential proteins. Microbial Cell Factories 11, 110 (2012).

35. Steel, H., Habgood, R., Kelly, C. L. & Papachristodoulou, A. In situ characterisation and manipulation of biological systems with Chi.Bio. PLOS Biology 18, e3000794 (2020).

36. Karst, S. M. et al. High-accuracy long-read amplicon sequences using unique molecular identifiers with Nanopore or PacBio sequencing. Nature Methods 18, 165– 169 (2021).

37. Henzler-Wildman, K. A. et al. A hierarchy of timescales in protein dynamics is linked to enzyme catalysis. Nature 450, 913–916 (2007).

38. Kerns, S. J. et al. The energy landscape of adenylate kinase during catalysis. Nature Structural & Molecular Biology 22, 124–131 (2015).

39. Kim, J. et al. Hidden resources in the Escherichia coli genome restore PLP synthesis and robust growth after deletion of the essential gene pdxB. Proceedings of the National Academy of Sciences 116, 24164–24173 (2019).

40. Peña, M. I., Davlieva, M., Bennett, M. R., Olson, J. S. & Shamoo, Y. Evolutionary fates within a microbial population highlight an essential role for protein folding during natural selection. Molecular Systems Biology 6, 387 (2010).

41. Arnold, F. H. Directed Evolution: Bringing New Chemistry to Life. Angewandte Chemie International Edition 57, 4143–4148 (2018).

42. Payne, J. L. & Wagner, A. The causes of evolvability and their evolution. Nature Reviews Genetics 20, 24–38 (2019).

43. Starr, T. N., Picton, L. K. & Thornton, J. W. Alternative evolutionary histories in the sequence space of an ancient protein. Nature 549, 409–413 (2017).

44. Xie, V. C., Pu, J., Metzger, B. P., Thornton, J. W. & Dickinson, B. C. Contingency and chance erase necessity in the experimental evolution of ancestral proteins. eLife 10, e67336 (2021).

45. Metzger, B. P. H., Park, Y., Starr, T. N. & Thornton, J. W. Epistasis facilitates functional evolution in an ancient transcription factor. eLife https://doi.org/10.7554/elife.88737.2 (2024) doi:10.7554/elife.88737.2.

46. Fröhlich, C. et al. Epistasis arises from shifting the rate-limiting step during enzyme evolution of a β-lactamase. Nature Catalysis 7, 499–509 (2024).

47. Bridgham, J. T., Ortlund, E. A. & Thornton, J. W. An epistatic ratchet constrains the direction of glucocorticoid receptor evolution. Nature 461, 515–519 (2009).

48. Akanuma, S. & Yamagishi, A. Biotechnology of Extremophiles:, Advances and Challenges. Grand Challenges in Biology and Biotechnology 581–596 (2016) doi:10.1007/978-3-319-13521-2_20.

49. Akanuma, S. et al. Experimental evidence for the thermophilicity of ancestral life. PNAS 110, 11067–11072 (2013).

50. Williams, P. D., Pollock, D. D., Blackburne, B. P. & Goldstein, R. A. Assessing the Accuracy of Ancestral Protein Reconstruction Methods. PLoS Computational Biology 2, e69 (2006).

51. Susko, E. & Roger, A. J. Problems With Estimation of Ancestral Frequencies Under Stationary Models. Systematic Biology 62, 330–338 (2013).

52. Sternke, M., Tripp, K. W. & Barrick, D. Chapter Seven The use of consensus sequence information to engineer stability and activity in proteins. Methods in Enzymology 643, 149–179 (2020).

53. McKinney, W. Data Structures for Statistical Computing in Python. in Proceedings of the 9th Python in Science Conference (eds Walt, S. van der & Millman, J.) 56–61 (2010). doi:10.25080/Majora-92bf1922-00a.

54. Harris, C. R. et al. Array programming with NumPy. Nature 585, 357–362 (2020).

55. Virtanen, P. et al. SciPy 1.0: fundamental algorithms for scientific computing in Python. Nature Methods 17, 261–272 (2020).

56. J. D. Hunter. Matplotlib: A 2D Graphics Environment. Computing in Science & Engineering 9, 90–95 (2007).

57. Abramson, J. et al. Accurate structure prediction of biomolecular interactions with AlphaFold 3. Nature 630, 493–500 (2024).

58. Goddard, T. D. et al. UCSF ChimeraX: Meeting modern challenges in visualization and analysis. Protein Science 27, 14–25 (2018).

59. Hanahan, D. Studies on transformation of Escherichia coli with plasmids. Journal of molecular biology 166, 557–580 (1983).

60. Datsenko, K. A. & Wanner, B. L. One-step inactivation of chromosomal genes in Escherichia coli K-12 using PCR products. Proceedings of the National Academy of Sciences 97, 6640–6645 (2000).

61. Newville, M. et al. LMFIT: Non-Linear Least-Squares Minimization and Curve-Fitting for Python. https://doi.org/10.5281/zenodo.598352 doi:10.5281/zenodo.598352.

62. Katoh, K. & Toh, H. Recent developments in the MAFFT multiple sequence alignment program. Briefings in Bioinformatics 9, 286–298 (2008).

63. Cock, P. J. A. et al. Biopython: freely available Python tools for computational molecular biology and bioinformatics. Bioinformatics 25, 1422–1423 (2009).

64. Bachmann, M. rapidfuzz/RapidFuzz: Release 3.8.1. Zenodo 10.5281/zenodo.10938887 (2024).

65. Keeble, A. H. et al. DogCatcher allows loop-friendly protein-protein ligation. Cell Chemical Biology 29, 339–350.e10 (2022).

