## Supplementary Information for "Comparing the evolvability of an ancestrally reconstructed and modern adenylate kinase"

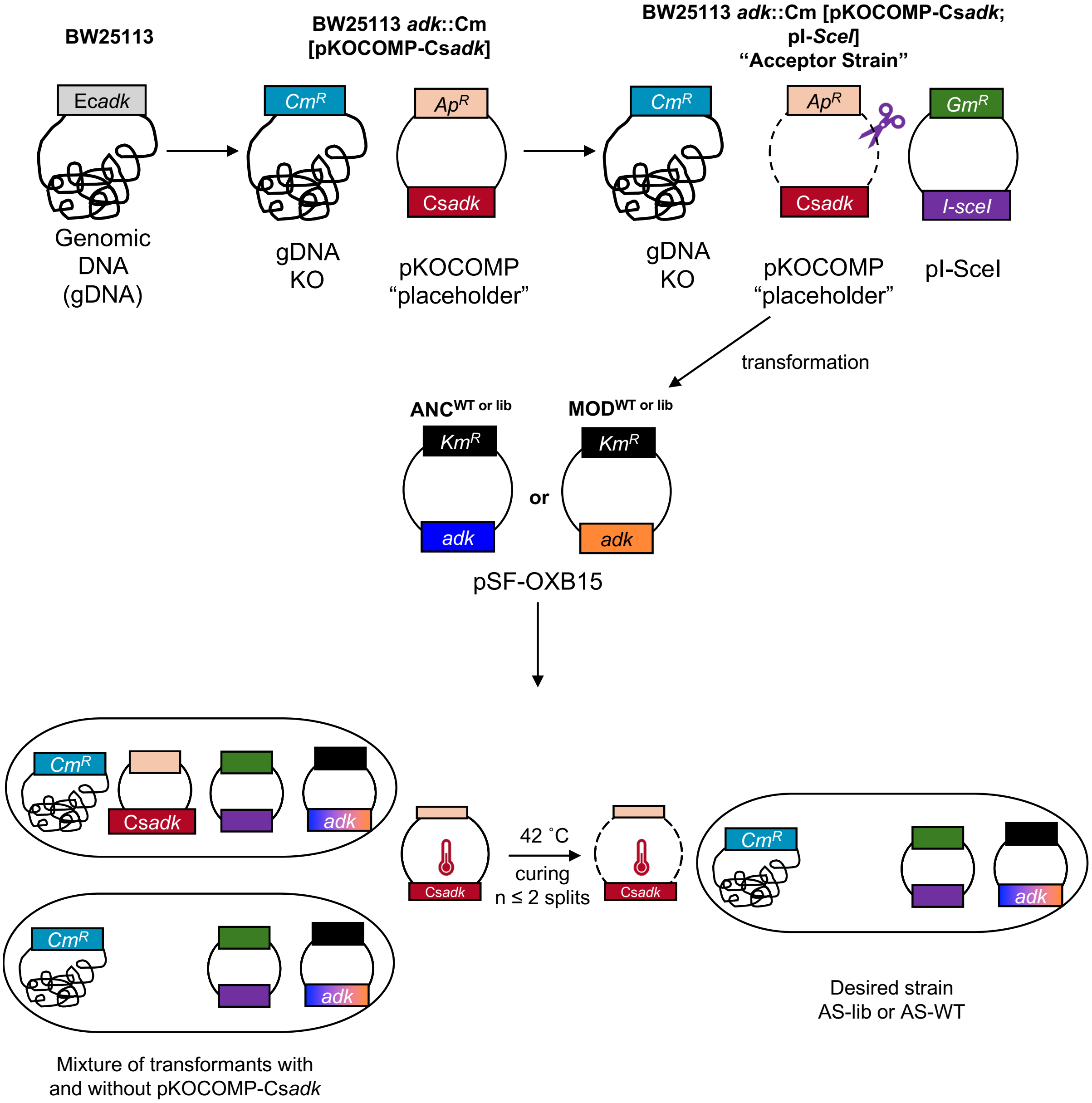


**Supplementary Fig. 1: The essential gene adenylate kinase can be knocked out, and "knocked in" to *E. coli* using a genetic replacement strategy.** *adk* knockout *E. coli* (BW25113) rescue strains were prepared by adapting a protocol from Billerbeck & Panke[1]. Genomic *adk* is replaced with a chloramphenicol resistance cassette using a λ-red recombination plasmid pKOCOMP that harbors Cs*adk*. The resulting strain was further transformed with pI-SceI harboring *I-sceI*, a restriction enzyme used to deplete pKOCOMP-Cs*adk* prior to transformation with pSFOXB15-*adk* (WT or library) of interest. To ensure complete curing from pKOCOMP-*Csadk*, thus full dependence of the cellular survival on pSFOXB15-*adk* encoded Adk activity, the strain was grown at 42 ºC utilizing the heat-sensitive origin of replication encoded on pKOCOMP. Abbreviations: Ec*adk*: genomic *E. coli* (BW25113) *adk*; $\text{Cm}^R$: chloramphenicol resistance gene; $\text{Ap}^R$: ampicillin resistance gene; $\text{Gm}^R$: gentamicin resistance gene; $\text{Km}^R$: kanamycin resistance gene; Cs*adk*: *adk* from *Caldanaerobacter subterraneus*; *I-SceI*: I-*Sce*I endonuclease.

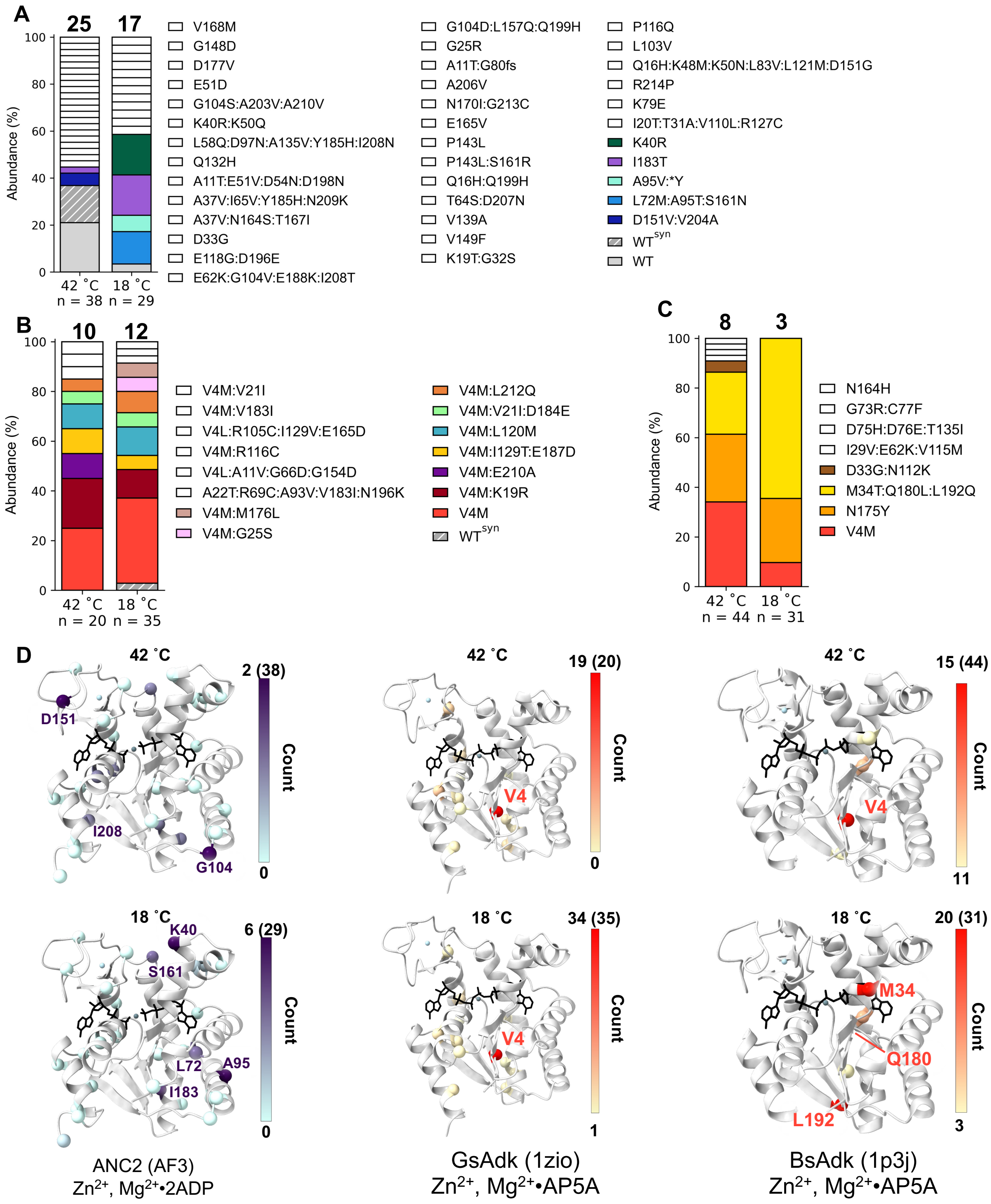


**Supplementary Fig. 2: ANC2 is more tolerant to mutations than Adks from *G. subterraneus and B. subtilis* during flask selection at 18 ºC. A-C** Population diversities (assessed by Sanger sequencing) before (42 ºC) and after flask selection at 18 ºC of ANC2 (**A**), GsAdklib (**B**), and BsAdklib (**C**) libraries show a larger population diversity in ANC2 than the extant enzymes GsAdk and BsAdk. The numbers of sequenced and unique clones are shown below and above the bars, respectively (WT$^{syn}$ = Adk with synonymous mutations, *Y = mutation of the stop codon to tyrosine). **D** Mutations are visualized on AlphaFold 3 predictions or crystal structures of Adks (grey cartoon, depicted as in **Fig. 1E**) as spheres colored according to abundance (see **A-C**).

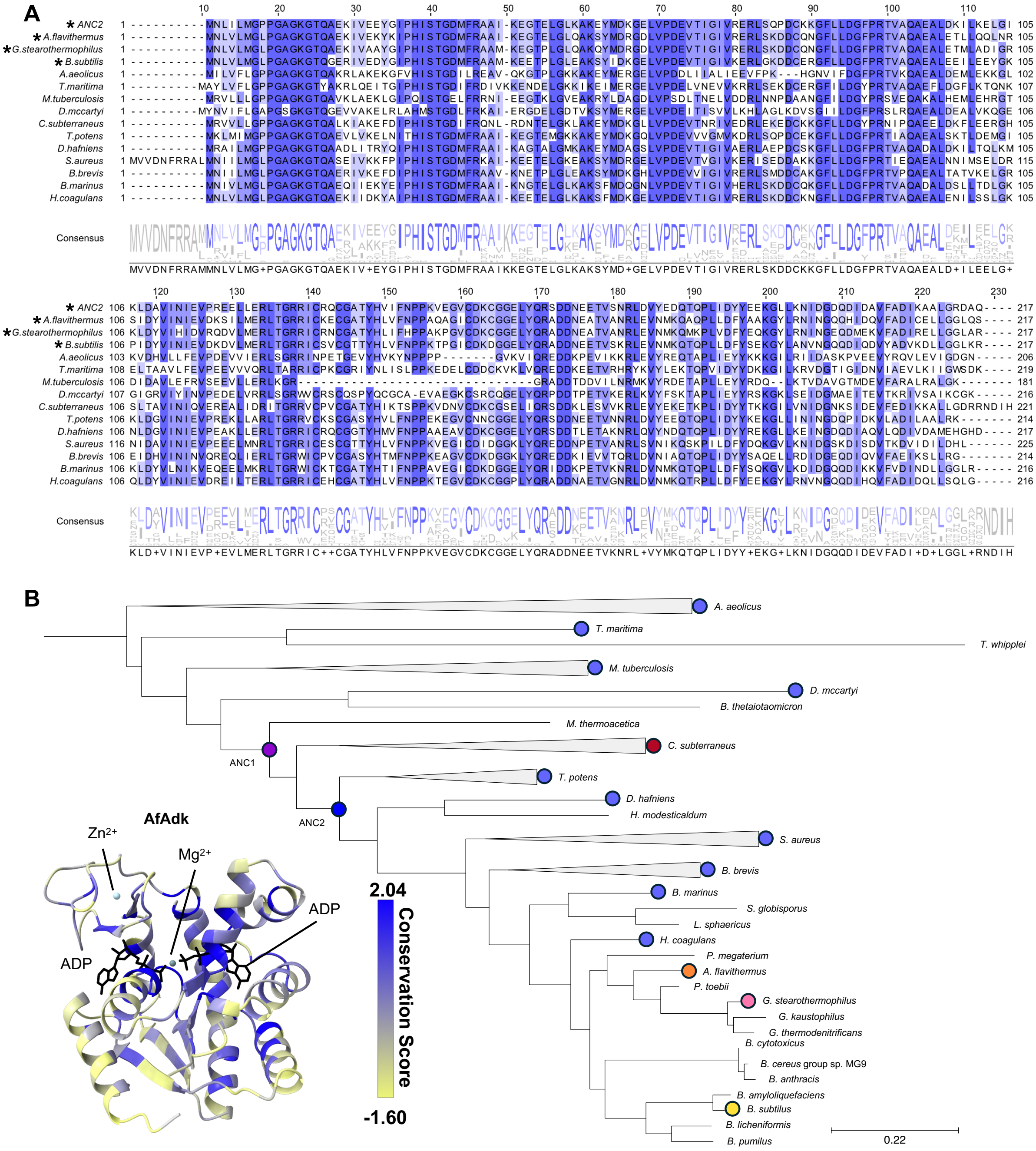


**Supplementary Fig. 3: Differences in Adk sequences occur globally throughout the enzyme's structure. A** Multiple sequence alignment of selected Adks from the phylogenetic tree of Nguyen *et al.*[2]. Adks used in selection experiments are labeled with an asterisk. Consensus sequence is displayed as a logo below the alignment. **B** AlphaFold 3 prediction of AfAdk (depicted as in **Fig. 1E**) colored by sequence conservation score (low: yellow & high: dark blue) of all Adk sequences found in the phylogenetic tree (**B**). Species used in (**A**) are labeled by circles and their scientific names. Ancestral Adk and modern Adks used in this study are colored in purple and dark blue as well as dark red, orange, pink, and yellow, respectively.

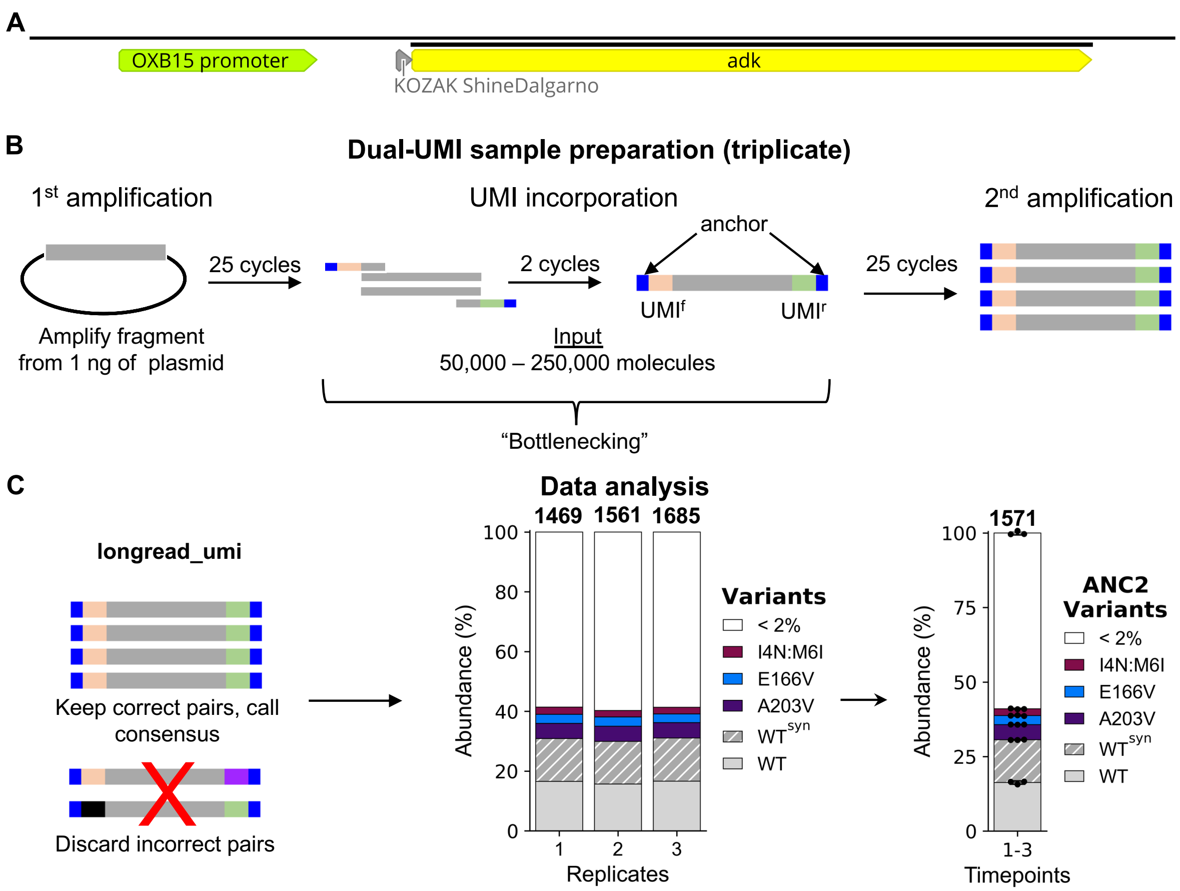


**Supplementary Fig. 4: High-sensitivity population diversity quantification using nanopore sequencing. A** Annotated fragment (including OXB15 promoter, 5' UTR and *adk*) amplified from selection timepoints for nanopore sequencing (image exported from Geneious version 2023.2.1). **B** Scheme of amplicon preparation and barcoding (adapted from Karst *et al.*[3]) to achieve robust quantification of individual variants present in the population. **C** Sample preparation and sequencing data (nanopore) analysis scheme. Nanopore sequencing was performed by Plasmidsaurus. Data analysis was performed utilizing the longread_umi pipeline[3] identifying consensus sequences by matching unique molecular identifier (UMI) pairs. This allows quantification of population diversity by calculating abundance from the number of UMIs found for each unique Adk variant (mean ± s.d. of n = 3). See Methods for more details.

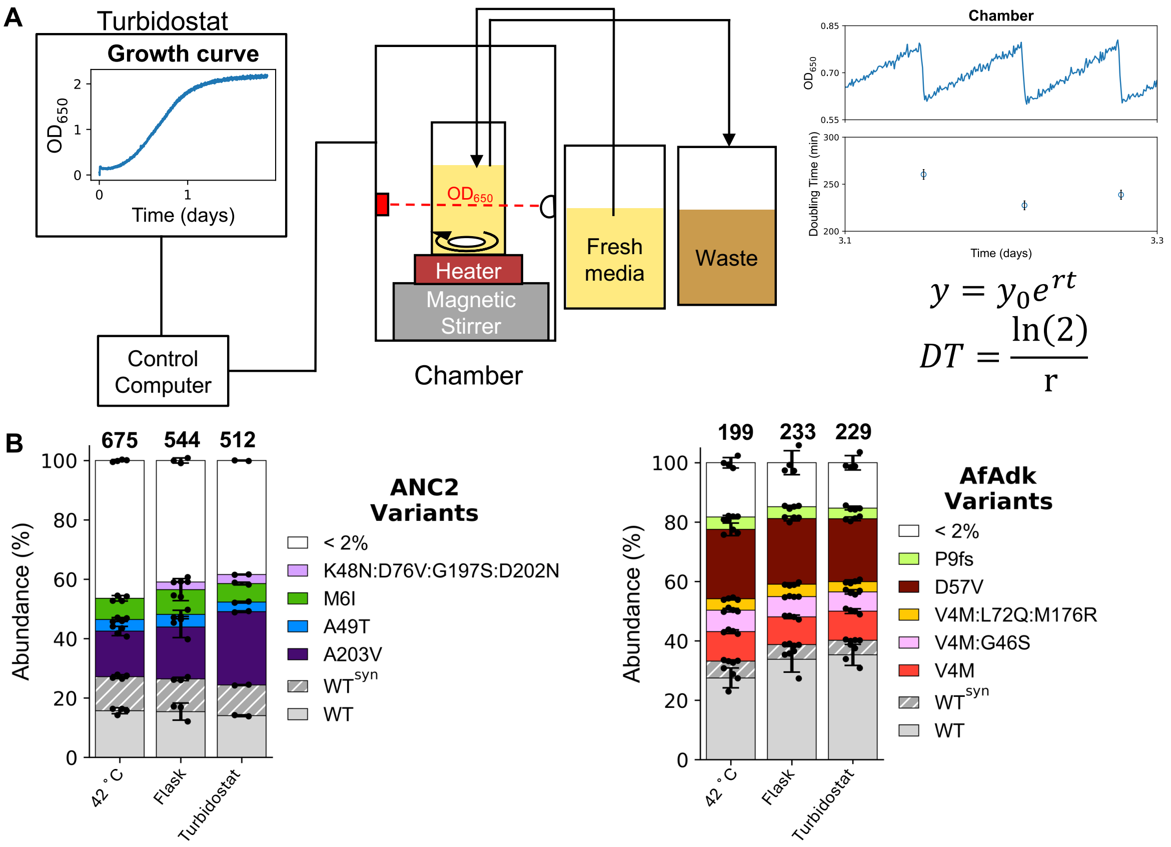


**Supplementary Fig. 5: Chi.Bio turbidostat is an automated alternative to labor-intensive log-phase selections in culture flasks. A** Abbreviated scheme of the Chi.Bio turbidostat[4]. Cultures were automatically maintained in log phase keeping $OD_{650}$ in a set range (0.6-0.8). The interval between dilutions was fit to a single exponential to calculate doubling time. **B** Turbidostat and culture flask selections are comparable: Abundance of ANC2 and AfAdk variants quantified from nanopore sequencing data (mean ± s.d. of n ≥ 2, unique variants shown above bars) at the following timepoints: 42 ºC, 20 ºC for 1 doubling time in a baffled culture flask, and ~ 24 h in the turbidostat at 20 ºC. $WT^{syn}$ = Adk with synonymous mutations, fs = frame shift. Data for each replicate are provided in Supplementary Tables 2 and 3.

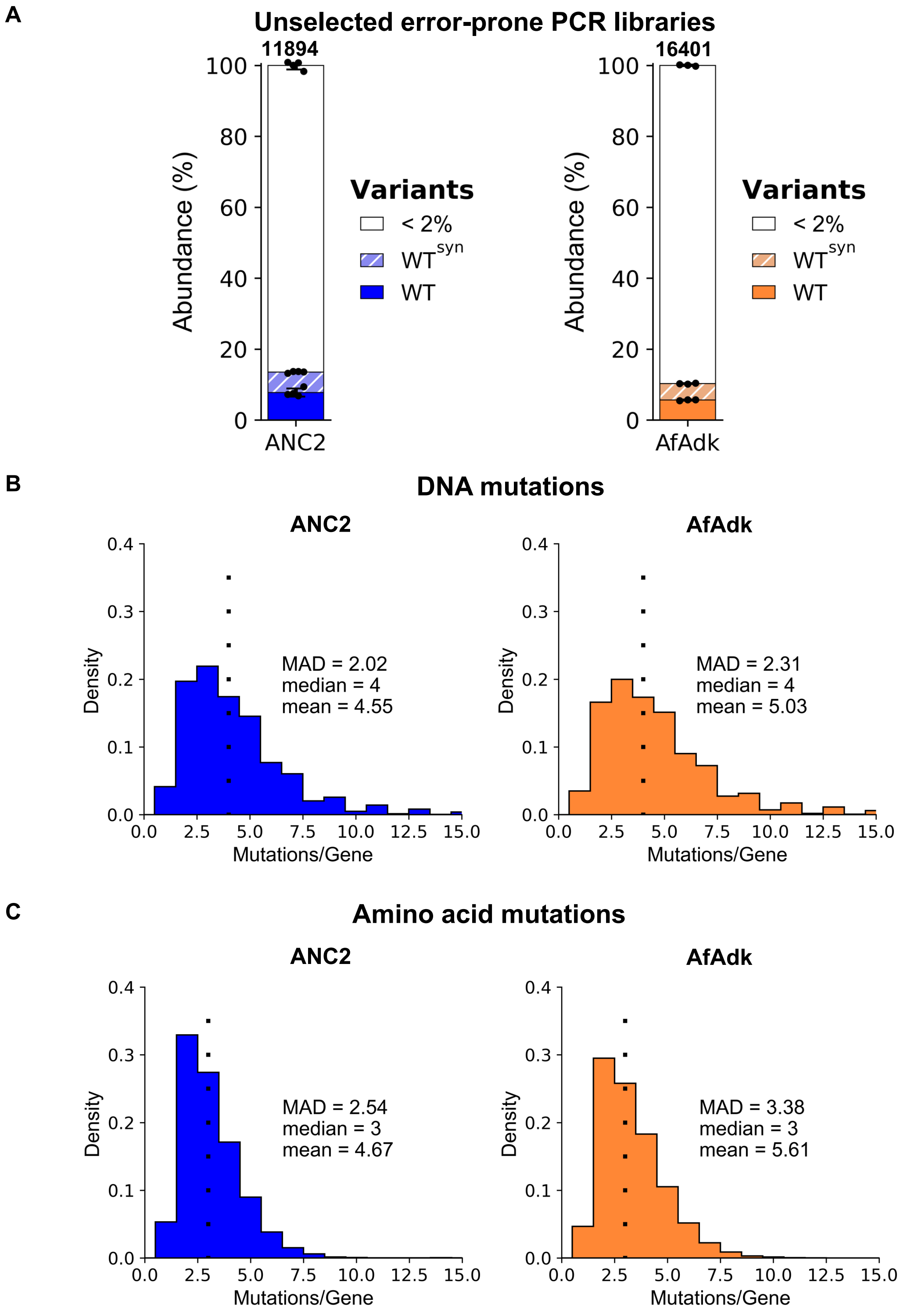


**Supplementary Fig. 6: The error-prone PCR library of AfAdk (AfAdklib) is slightly more diverse than ANC2lib. A** Starting library diversity quantified via nanopore sequencing (Supplementary Fig. 4) for ANC2lib and AfAdklib (mean ± s.d. of n ≥ 3; average number of unique variants indicated above bars). Data for each replicate are provided in Supplementary Tables 2 and 3. **B-C** Histograms of DNA mutations/gene for ANC2 and AfAdk library variants (**B**) and nonsynonymous (amino acid) mutations/gene for ANC2 and AfAdk library variants (**C**) show larger genetic diversity for AfAdk. MAD = mean absolute deviation around the median.

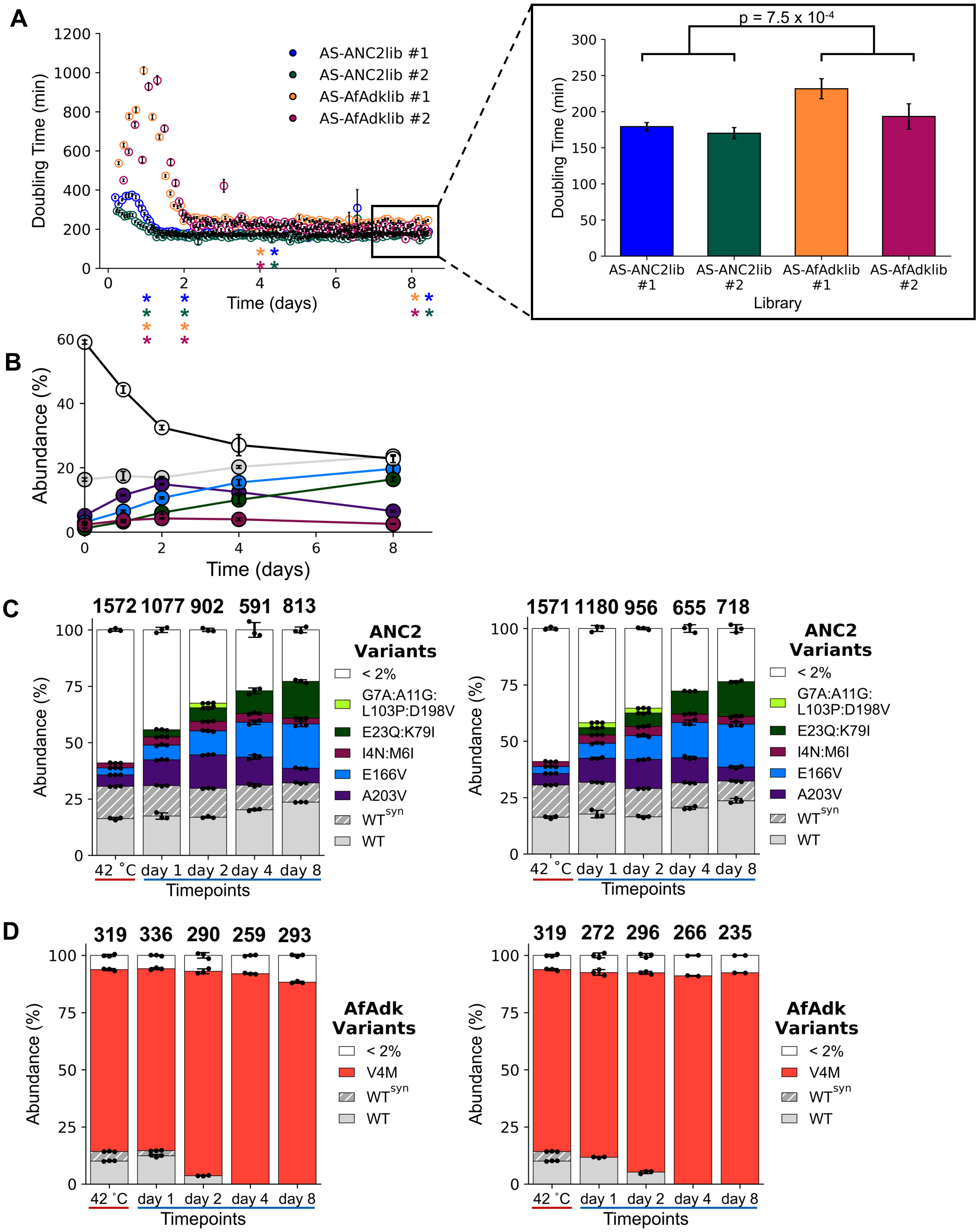


**Supplementary Fig. 7: ANC2 library strain displayed greater fitness and population diversity than the AfAdk library strain. A** Doubling times of AS-ANC2lib and AS-AfAdklib during turbidostat selection at 20 ºC (technical culture replicates n = 2). ANC2 library is significantly faster than AfAdk library after stabilization at selection day 8. (inset; Two-sided Kolmogorov-Smirnov test, $p = 7.5 \times 10^{-4}$). Colored asterisks denote timepoints at which sequencing samples were taken (sequencing data in (**C**) and (**D**)). **B** Plot of ANC2 variant enrichment for the highest abundance mutants (shown in (**C**), same colors are used) during 20 ºC selection. **C**-**D** Nanopore sequencing data for the two technical replicates of AS-ANC2lib (**C**) and AfAdklib (**D**) before (42 ºC) and throughout 20 ºC selection (mean ± s.d. of n ≥ 2; average number of unique variants shown above bars). 42 ºC refers to the sample taken prior to inoculation of turbidostat chambers (see plasmid curing, Supplementary Fig. 1). $WT^{syn}$ = Adk with synonymous mutations. Data for each replicate are provided in Supplementary Tables 2 and 3.

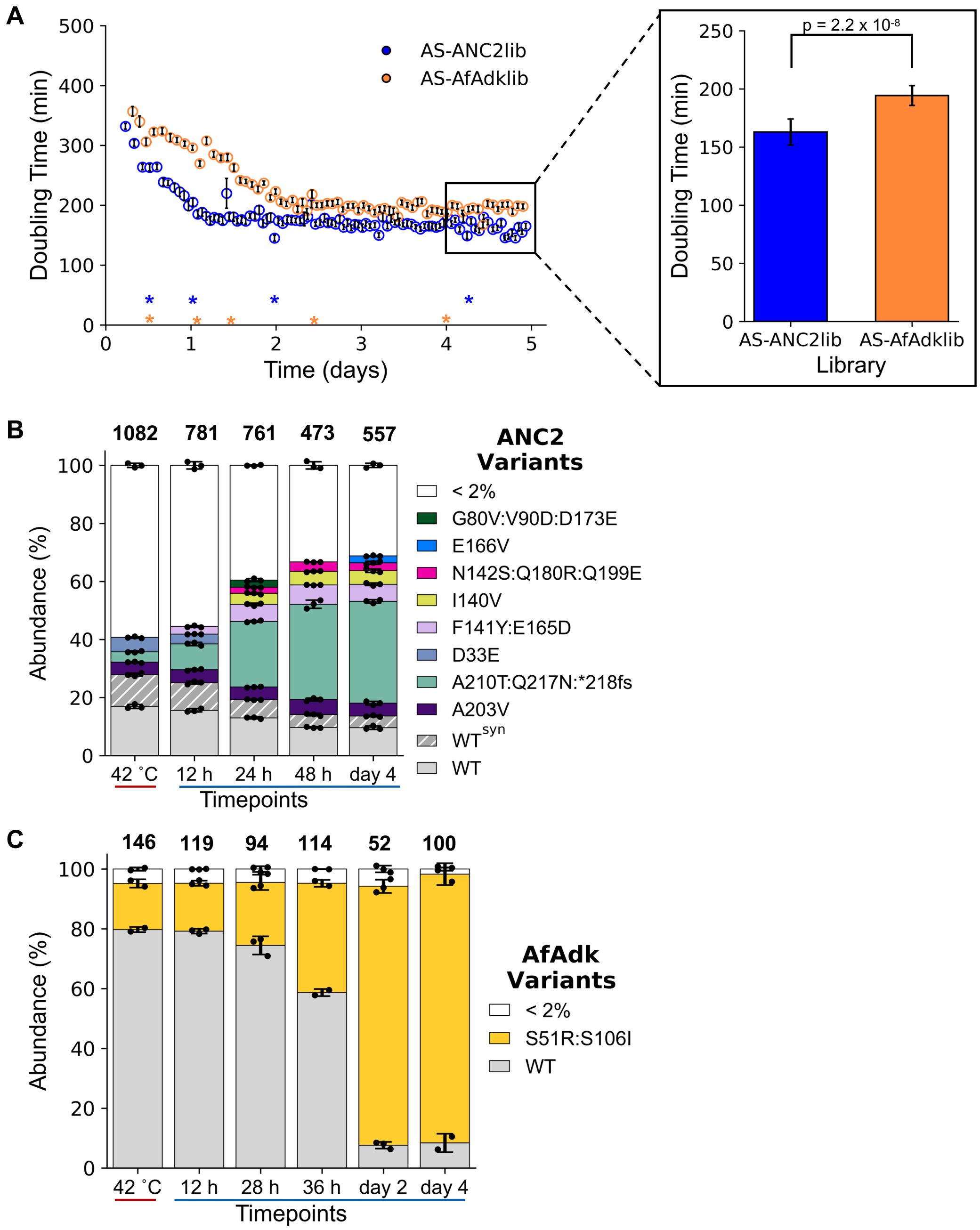


**Supplementary Fig. 8: AS-ANC2lib was reproducibly more fit and diverse than AS-AfAdklib. A** Doubling times of a biological replicate of AS-ANC2lib and AS-AfAdklib during selection at 20 ºC (compare to experiment Supplementary Fig. 7). Significant differences in stabilized doubling times after 8 days are shown in the inset (Two-sided Kolmogorov-Smirnov test, $p = 2.2 \times 10^{-8}$). Colored asterisks denote timepoints at which sequencing samples were taken (data in (**B**) and (**C**)). **B**-**C** Stacked bar graphs depicting AS-ANC2lib (**B**) and AfAdklib (**C**) population diversity (mean ± s.d. of n ≥ 2; average number of unique variants shown above bars). 42 ºC refers to the sample taken prior to inoculation of turbidostat chambers (see plasmid curing, Supplementary Fig. 1). $WT^{syn}$ = Adk with synonymous mutations, *fs = a frame shift at the stop codon. Data for each replicate are provided in Supplementary Tables 2 and 3.

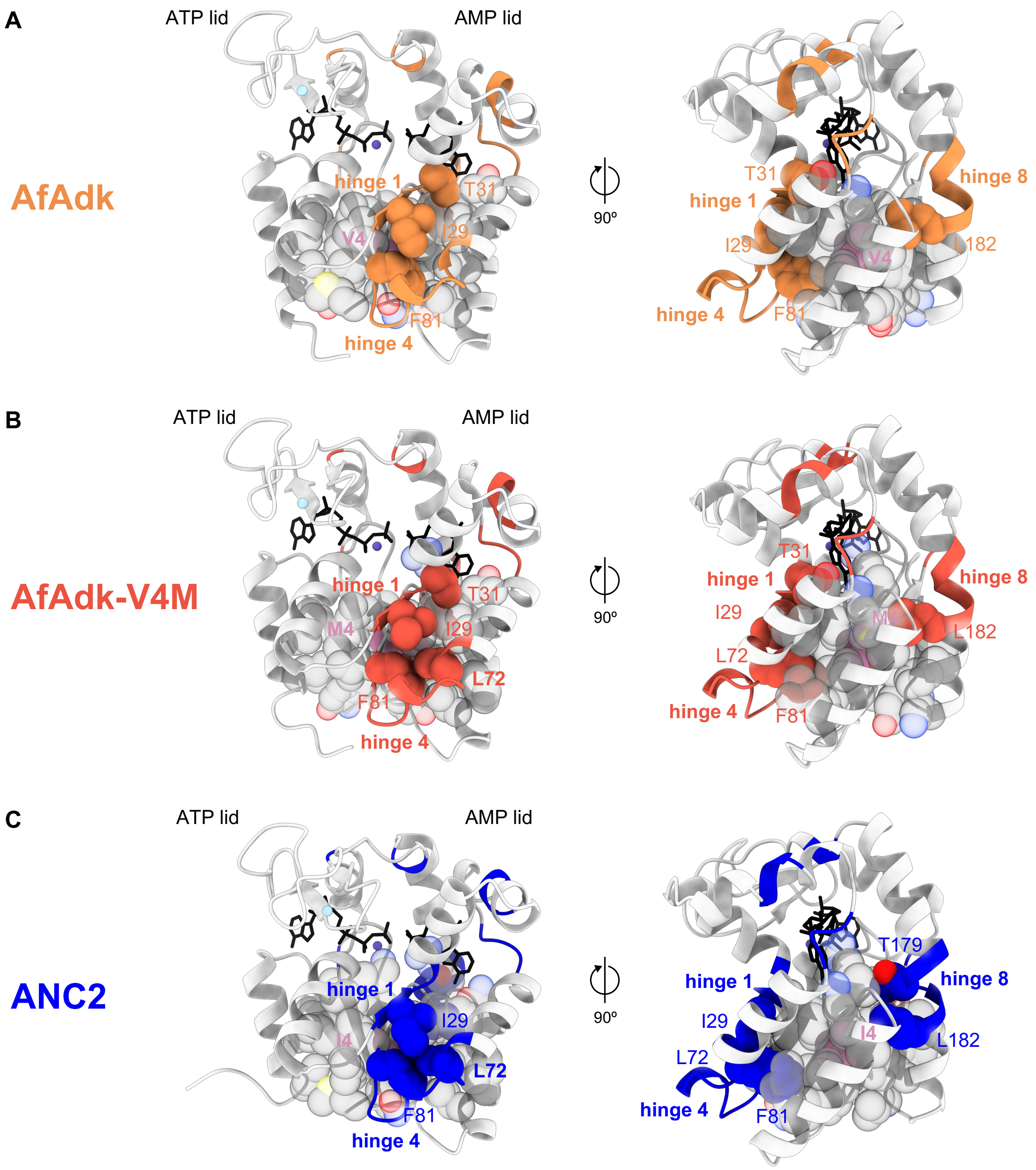


**Supplementary Fig. 9: Differences in packing of hinge 4 might alter kinetics of AfAdk-V4M. A-C** AlphaFold 3 predictions (grey cartoon; depicted as in **Fig. 1E**) of AfAdk (**A**), AfAdk-V4M (**B**), and ANC2 (**C**). All eight hinge regions (colored cartoon) as defined by Henzler-Wildman *et al.*[5] involved in rate-limiting opening/closing dynamics that could possibly be affected by the mutation at position 4 are highlighted accordingly. 1st and 2nd shell residues of residue 4 (V4, M4 and I4, respectively; pink spheres) are depicted as gray transparent spheres, unless they are also part of hinges (non-transparent spheres colored according to hinge color). In AfAdk-V4M (**B**), L72 (hinge 4) lies in the 2nd contact shell of M4, while packing against 2nd shell hinge residues I29 (hinge 1) and F81 (hinge 4). Unlike the slower AfAdk WT (**A**), where L72 is placed outside of the 2nd shell of V4M, this arrangement in V4M mirrors the faster ANC2 WT (**C**), where L72 is part of I4's 2nd shell contact as well. Nitrogen, oxygen, phosphorus, sulfur, and carbon are colored blue, red, orange, yellow, and grey (unless highlighted otherwise), respectively.

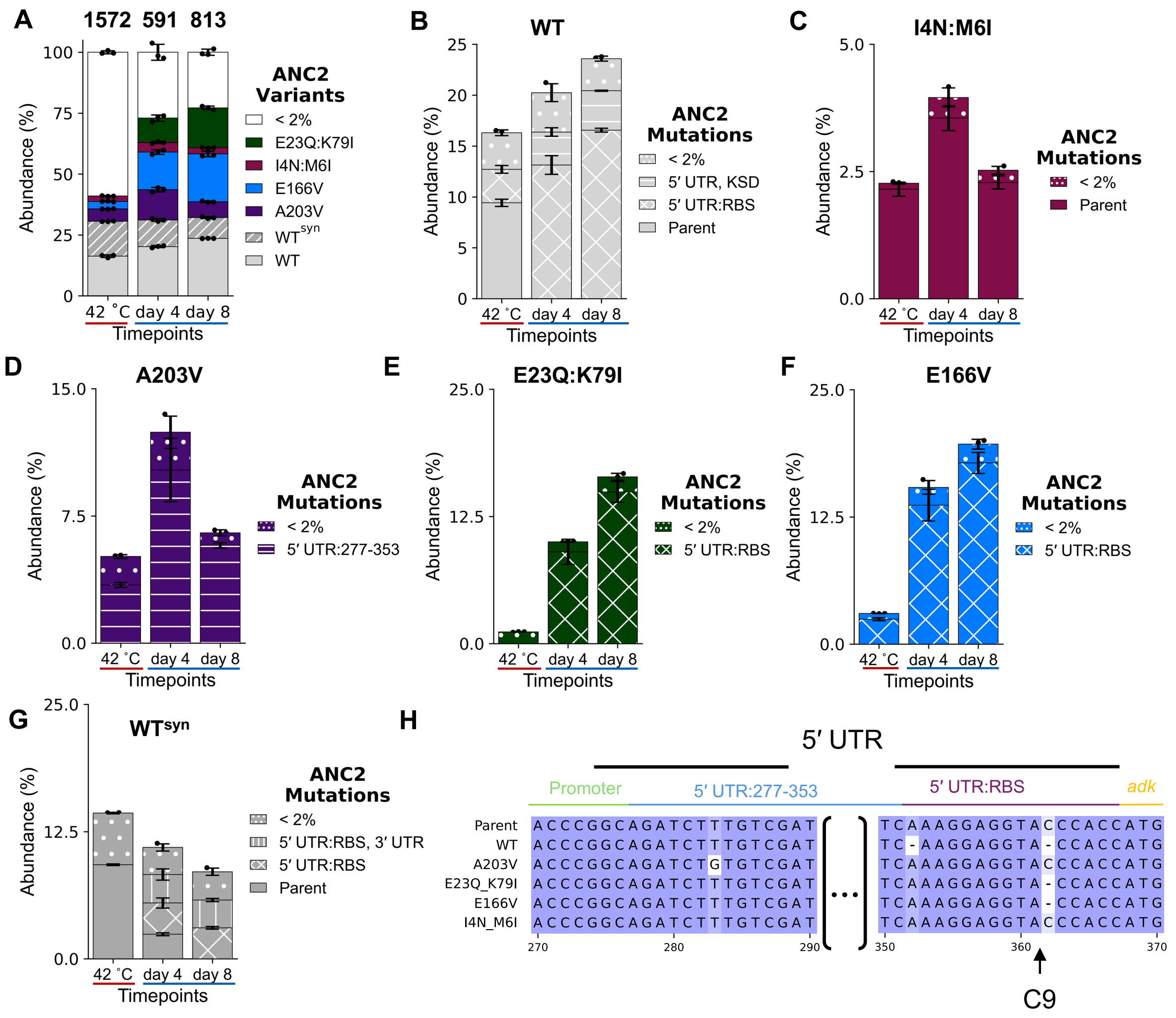


**Supplementary Fig. 10. Mutations in the 5′ UTR, particularly the RBS, increased fitness of ANC2.** **A-G** Stacked bar graphs highlight off-target mutation impact by depicting abundance before (42 ºC) and during selection at 20 ºC of AS-ANC2lib variants (**A**) and off-target mutations for ANC2 WT (**B**), I4N:M6I (**C**), A203V (**D**), E23Q:K79I (**E**), E166V (**F**), and WT$^{syn}$ (**G**) prior (42 ºC) to and throughout selection at 20 ºC (mean ± s.d. of n = 3). **H** High-abundance variants' 5′ UTR (277-353 region and RBS) are shown aligned to the parent vector sequence. The WT sequence displayed is enriched ANC2 WT (**B**). 5′ UTR (including *adk* start codon ATG: 368-370). WT$^{syn}$ is not included in the alignment due to the heterogeneity of synonymous *adk* off-target mutations.

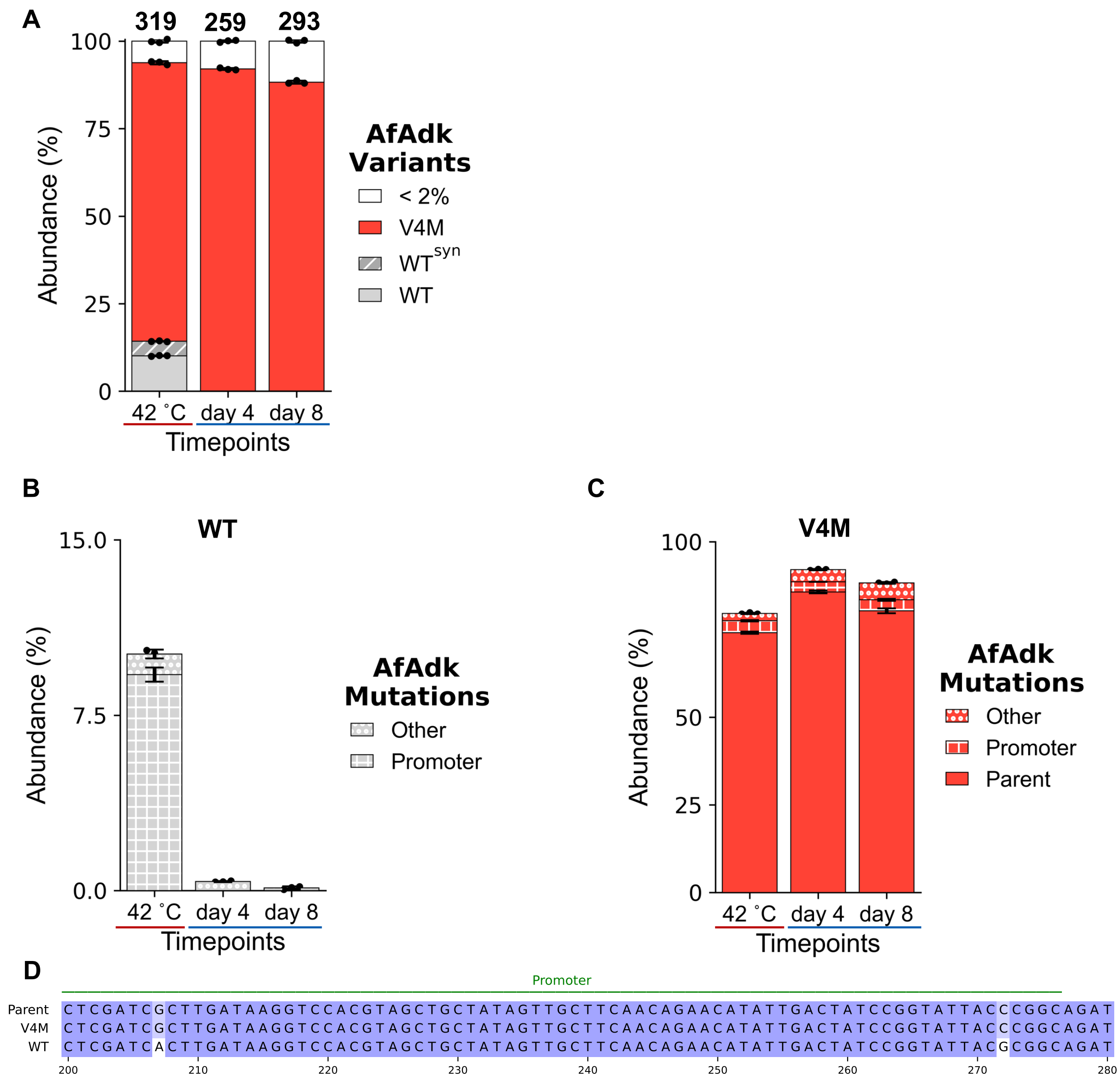


**Supplementary Fig. 11: AfAdk-V4M outcompeted AfAdk WT containing two promoter mutations. A-C** Stacked bar graphs showing population diversity prior to (42 ºC) and throughout selection at 20 ºC for AS-AfAdklib variants (**A**) and off-target mutations for depleting AfAdk WT (**B**) and enriching V4M (**C**) (mean ± s.d. of n = 3). **D** AfAdk WT and AfAdk-V4M promoter regions aligned to the parent OXB-15 sequence.

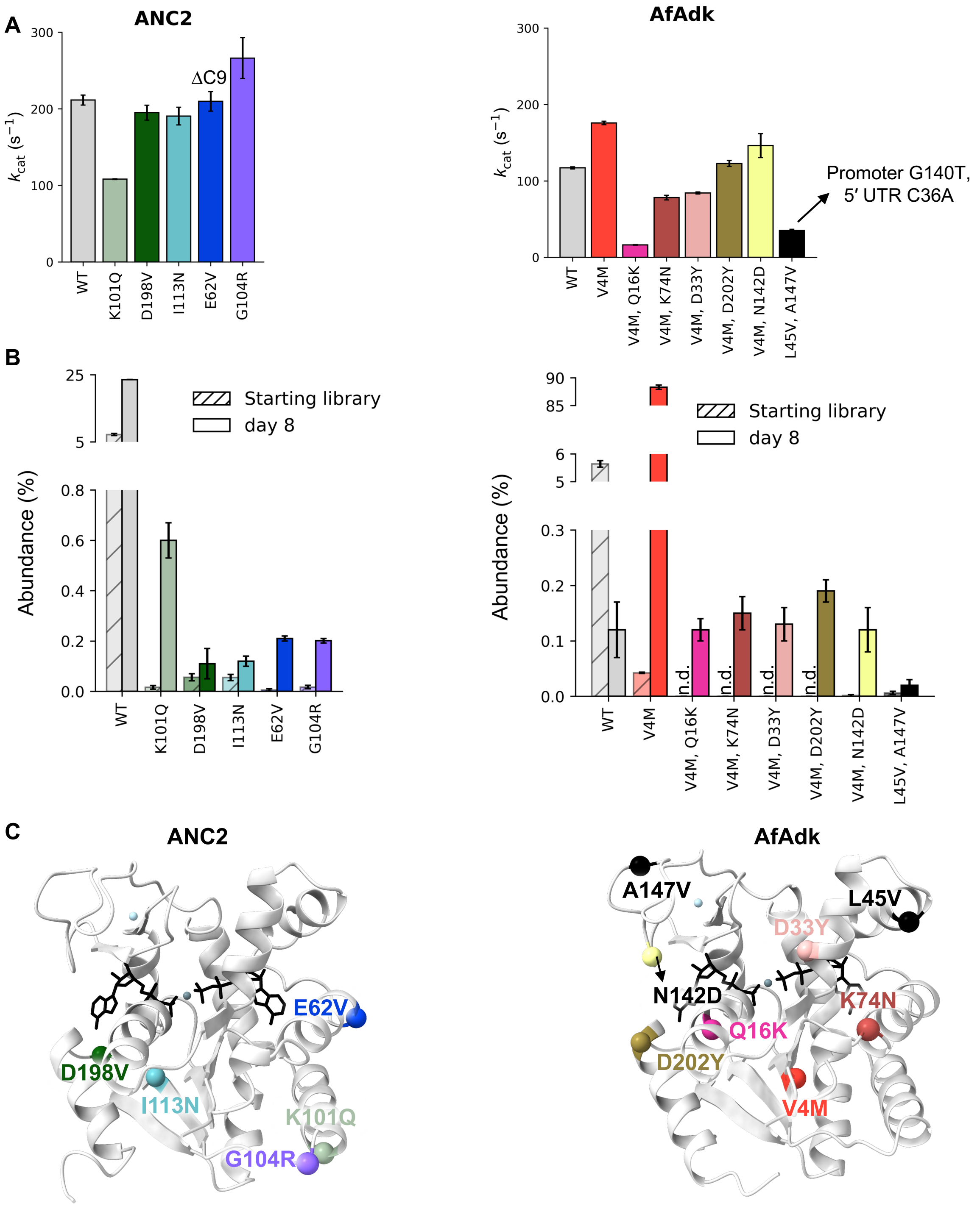


**Supplementary Fig. 12: Most low-abundance Adk variants after selection had similar $k_{cat}$ values to WT. A** $k_{cat}$ values of ATP production by low-abundance Adks (ANC2: left & AfAdk: right) variants measured at 20 ºC, via coupled assay (mean ± s.d. of n = 3 replicates). Off-target mutations, if present, are indicated above bars (see Supplementary Fig. 10H). **B** Abundance of each variant quantified by nanopore sequencing (ANC2: left & AfAdk: right) in the starting library and after 8 days of selection at 20 ºC (mean ± s.d. of n ≥ 3; n.d. = not detected). **C** Mutations are globally distributed as depicted by colored spheres onto AlphaFold 3 predictions of Adks (depicted as in **Fig. 1E**).

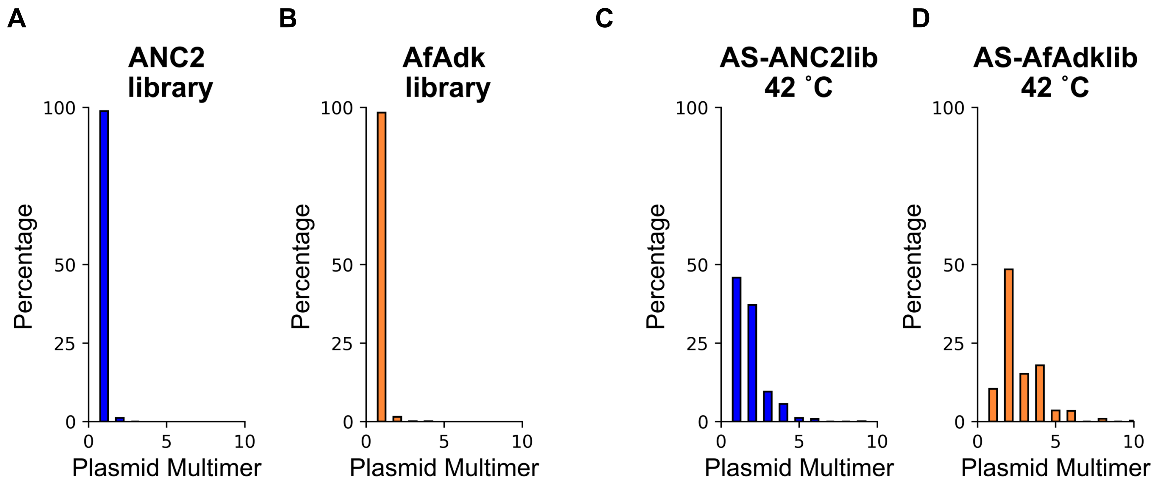


**Supplementary Fig. 13: Plasmid multimerization was found in all biological samples and potentially contributed to fitness in *in vivo* selection at 20 ºC. A-D** Histograms representing the size of all plasmids miniprepped from starting libraries ANC2lib (**A**) and AfAdklib (**B**) as well as the acceptor strains (AS) transformed with these staring libraries: ANC2lib (**C**) and AfAdklib (**D**) after plasmid (placeholder) curing at 42 ºC.

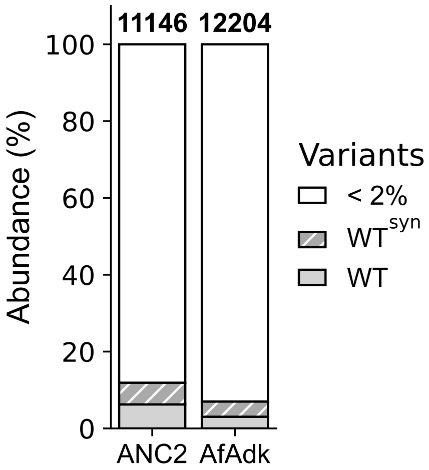


**Supplementary Fig. 14: Error-prone PCR libraries used for *in vitro* selection are similarly diverse to those used for *in vivo* selection (see Supplementary Fig. 6A).** Population diversity quantified by nanopore sequencing of *in vitro* epPCR libraries of ANC2 or AfAdk.

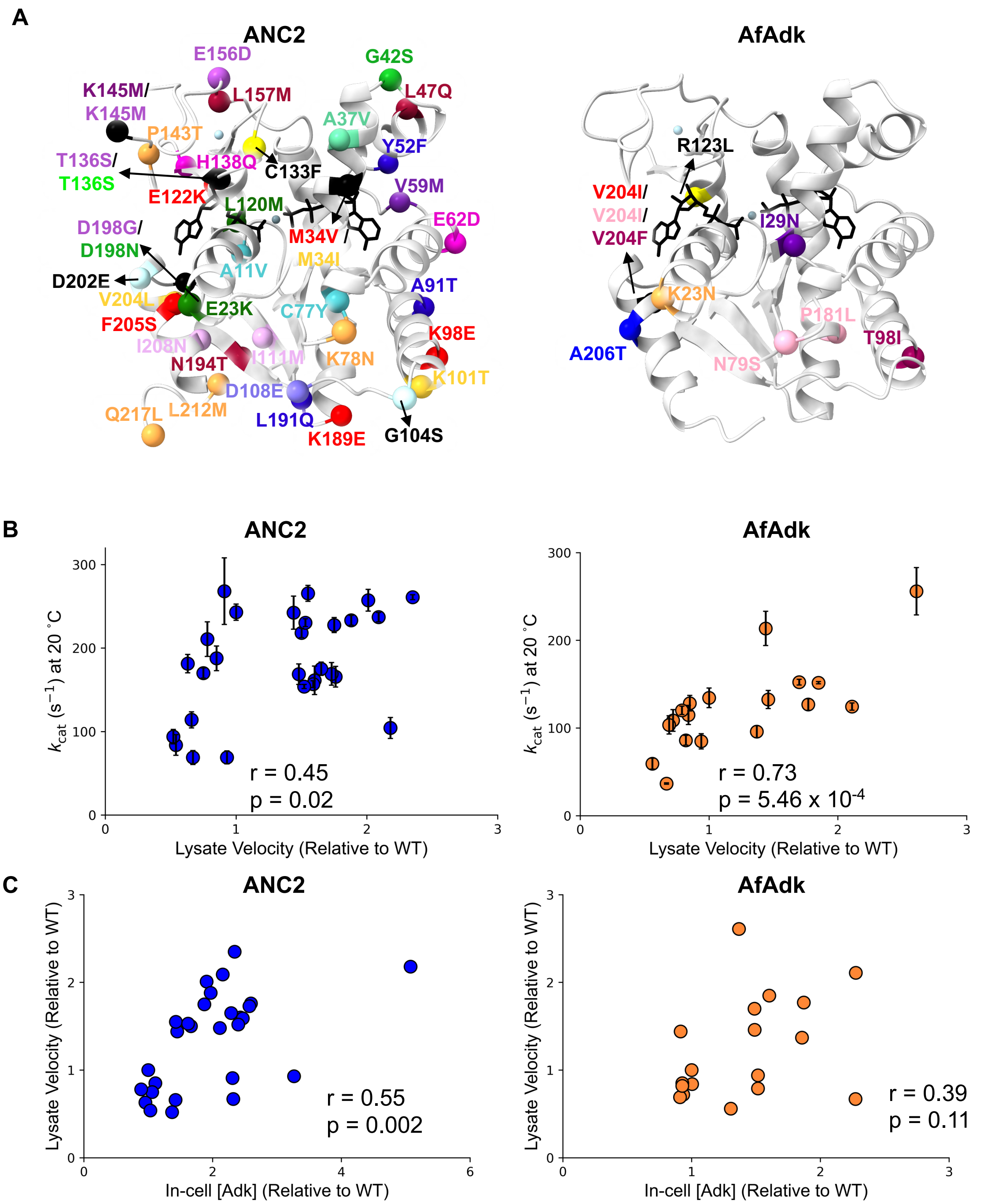


**Supplementary Fig. 15: ANC2 variants increase both activity and soluble expression *in vitro*. A** Globally distributed mutations identified during *in vitro* selections at 20 ºC are colored by variant (**Fig. 4B**, C) and plotted onto AlphaFold 3 predictions of ANC2 or AfAdk (grey cartoon; depicted as in **Fig. 1E**). Positions mutated multiple times are represented as black spheres with an arrow pointing to the mutation(s) found at that position, colored by variant (see Fig. 4B-C). **B-C** Lysate velocity of ANC2 variants correlated more with cellular Adk concentration than $k_{cat}$ (left), whereas AfAdk lysate activity correlated much more with $k_{cat}$ than Adk concentration (right). This reflects the differences in catalytic improvement between ANC2 and AfAdk (see Fig. 4B). **B** Correlation of $k_{cat}$ values plotted against lysate velocity for ANC2 (left) and AfAdk (right) variants (Pearson $r_{ANC2}$ = 0.45, $p_{ANC2}$ = 0.02; Pearson $r_{AfAdk}$ = 0.73, $p_{AfAdk}$ = 5.46 x $10^{-4}$). **C** Correlation of lysate velocities of ANC2 and AfAdk variants plotted against Adk concentration in the lysate for ANC2 (left, Pearson r = 0.55, p = 0.002), but not for AfAdk variants (right, Pearson r = 0.39, p = 0.11).

**Supplementary Table 1: Primers.**

| Name | Sequence (5′ - 3′ ) | Notes |
|---|---|---|
| pSFOXB15_epPCR_for | GCTCTCGAATTCAAAGGAGGTACCCACC | |
| pSFOXB15_epPCR_rev | GTCAGTCAGTGCAGGAGGAGACAACTTCTAG | |
| pET17b_epPCR_for | GTTTAACTTTAAGAAGGAGATATACAT | |
| pET17b_ANC2_epPCR_rev | CCCTGGAAATACAGATTCTC | |
| pET17b_Af_epPCR_rev | ACCTTGAAAATACAGGTTTTC | |
| pSFOXB15_ont_for | GTTATGCTATCAATCGTTGC | |
| pSFOXB15_ont_rev | GAAGGTACGCTGTATCTCAG | |
| ont_ss_dumi_for | CAAGCAGAAGACGGCATACGAGATNNNYRNNNYRNNNYRNNNGTTATGCTATCAATCGTTGC | Adapted from Karst *et al.*[3] for complementation to pSF-OXB15 fragment |
| ont_ss_dumi_rev | AATGATACGGCGACCACCGAGATCNNNYRNNNYRNNNYRNNNGAAGGTACGCTGTATCTCAG | Adapted from Karst *et al.*[3] for complementation to pSF-OXB15 fragment |
| ont_synthetic_primer_for | CAAGCAGAAGACGGCATACGAGAT | Karst *et al.*[3] |
| ont_synthetic_primer_rev | AATGATACGGCGACCACCGAGATC | Karst *et al.*[3] |

Primers used to make error-prone PCR libraries and to prepare samples for nanopore sequencing.

**Supplementary Table 2: Nanopore sequencing data of AS-ANC2lib replicates.**

| Experiment | Fig. | Timepoint | # vars 1 | # vars 2 | # vars 3 | # vars 4 | mean | s.d. |
|---|---|---|---|---|---|---|---|---|
| N/A | S6 | Starting library | 7714 | 14511 | 12338 | 13012 | 11894 | 2538 |
| 20230427 | S5 | 42 ˚C | 661 | 767 | 657 | 616 | 675 | 56 |
| 20230427 | S5 | sf | 586 | 526 | 519 | | 544 | 30 |
| 20230427 | S5 | T | 511 | 512 | | | 512 | 1 |
| 20240218 | 2, S7 | 42 ˚C | 1469 | 1561 | 1685 | | 1572 | 89 |
| 20240218 | 2, S7 | 24 h (Rep 1) | 1247 | 996 | 989 | | 1077 | 120 |
| 20240218 | 2, S7 | 48 h (Rep 1) | 975 | 1018 | 713 | | 902 | 135 |
| 20240218 | S82, S7 | day 4 (Rep 1) | 758 | 468 | 547 | | 591 | 122 |
| 20240218 | S82, S7 | day 8 (Rep 1) | 991 | 763 | 685 | | 813 | 130 |
| 20240218 | 2, S7 | 24 h (Rep 2) | 936 | 1257 | 1347 | | 1180 | 176 |
| 20240218 | 2, S7 | 48 h (Rep 2) | 821 | 1016 | 1030 | | 956 | 95 |
| 20240218 | 2, S7 | day 4 (Rep 2) | 586 | 819 | 559 | | 655 | 117 |
| 20240218 | 2, S7 | day 8 (Rep 2) | 929 | 624 | 601 | | 718 | 149 |
| 20231126 | S8 | 42 ˚C | 1302 | 1045 | 898 | | 1082 | 167 |
| 20231126 | S8 | 12 h | 725 | 873 | 745 | | 781 | 66 |
| 20231126 | S8 | 24 h | 708 | 832 | 742 | | 761 | 52 |
| 20231126 | S8 | 48 h | 520 | 491 | 408 | | 473 | 47 |
| 20231126 | S8 | day 4 | 748 | 517 | 407 | | 557 | 142 |

Number of variants for each sequenced replicate from ANC2 library samples.

**Supplementary Table 3: Nanopore sequencing data of AS-AfAdklib replicates.**

| Experiment | Fig. | Timepoint | # vars 1 | # vars 2 | # vars 3 | # vars 4 | mean | s.d. |
|---|---|---|---|---|---|---|---|---|
| N/A | S6 | Starting library | 12057 | 18299 | 18846 | | 16401 | 3080 |
| 20230427 | S5 | 42 ˚C | 403 | 159 | 198 | 510 | 253 | 107 |
| 20230427 | S5 | sf | 200 | 338 | 207 | 526 | 318 | 132 |
| 20230427 | S5 | T | 226 | 286 | 237 | 511 | 250 | 26 |
| 20240218 | 2, S7 | 42 ˚C | 271 | 318 | 367 | | 319 | 39 |
| 20240218 | 2, S7 | 24 h (Rep 1) | 334 | 349 | 326 | | 336 | 10 |
| 20240218 | 2, S7 | 48 h (Rep 1) | 293 | 303 | 274 | | 290 | 12 |
| 20240218 | 2, S7 | day 4 (Rep 1) | 252 | 215 | 311 | | 259 | 40 |
| 20240218 | 2, S7 | day 8 (Rep 1) | 300 | 295 | 285 | | 293 | 6 |
| 20240218 | 2, S7 | 24 h (Rep 2) | 308 | 270 | 239 | | 272 | 28 |
| 20240218 | 2, S7 | 48 h (Rep 2) | 305 | 298 | 286 | | 296 | 8 |
| 20240218 | 2, S7 | day 4 (Rep 2) | 241 | 291 | | | 266 | 25 |
| 20240218 | 2, S7 | day 8 (Rep 2) | 257 | 213 | | | 235 | 22 |
| 20231126 | S8 | 42 ˚C | 160 | 131 | | | 146 | 15 |
| 20231126 | S8 | 12 h | 121 | 92 | 145 | | 119 | 22 |
| 20231126 | S8 | 28 h | 69 | 127 | 87 | | 94 | 24 |
| 20231126 | S8 | 36 h | 109 | 119 | | | 114 | 5 |
| 20231126 | S8 | day 2 | 60 | 48 | 48 | | 52 | 6 |
| 20231126 | S8 | day 4 | 54 | 146 | | | 100 | 46 |

Number of variants for each sequenced replicate from AfAdk library samples.

**Supplementary Table 4: Enzymatic activity at 20 ºC and thermostability data for ANC2 and AfAdk variants from *in vivo* selection.**

| Parent | name | $k_{cat}$ ($s^{-1}$) | $k_{cat}$ s.d. | $T_m$ (˚C) | $T_m$ s.d. | $k_{cat}/K_{m, ADP}$ ($M^{-1}s^{-1}$) | $k_{cat}/K_m$ error |
|---|---|---|---|---|---|---|---|
| ANC2 | WT | 245 | 13 | 79.7 | 0.5 | 2.4E+05 | 2.6E+04 |
| ANC2 | A203V | 212 | 13 | 78.5 | 0.3 | 2.5E+05 | 4.7E+03 |
| ANC2 | E166V | 258 | 21 | 80.1 | 0.3 | 2.5E+05 | 6.3E+03 |
| ANC2 | I4N, M6I | 135 | 4 | 67.9 | 0.1 | 1.1E+05 | 2.1E+03 |
| ANC2 | E23Q, K79I | 223 | 4 | 77.1 | 0.3 | 2.6E+05 | 1.2E+04 |
| AfAdk | WT | 136 | 2 | 71.4 | 0.2 | 3.1E+05 | 4.5E+04 |
| AfAdk | V4M | 152 | 5 | 65.8 | 0.6 | 2.6E+05 | 2.0E+04 |

Activity was measured via coupled assay in ATP direction with 5 mM $Mg^{2+}$•ADP (mean ± s.d. of n ≥ 3 replicates) and $T_m$ values were measured by a thermal shift assay (with SYPRO Orange) from 20 to 95 ºC.

**Supplementary Table 5: Activity and thermostability data for ANC2 variants selected in *in vitro* experiments.**

| ID | Variant | Synonymous mutations | Lysate activity (normalized) | $k_{cat}$ ($s^{-1}$) | $k_{cat}$ s.d. | $T_m$ (˚C) | $T_m$ s.d. | Cellular [Adk], mM | Background (vector) mutations? |
|---|---|---|---|---|---|---|---|---|---|
| N/A | WT | | 1 | 243 | 10 | 79.7 | 0.5 | 6 | |
| Plate 5, H9 | C133F | | 2.18 | 104 | 13 | 78.4 | 0.1 | 29 | |
| Plate 6, D1 | A37V | L96L, T125T, I140I, D200D | 2.01 | 257 | 13 | 80 | 1 | 11 | |
| Plate 8, A12 | D108E | N2N | 1.53 | 230 | 5 | 76.4 | 0.7 | 9 | |
| Plate 8, G12 | V59M | | 1.48 | 169 | 12 | 77.7 | 0.3 | 12 | |
| Plate 5, H2 | A11V, C77Y | | 1.44 | 242 | 20 | 78.4 | 0.1 | 8 | |
| Plate 7, D10 | M34I, K101T, V204L | T64T, I195I | 1.52 | 154 | 3 | 77.9 | 0.6 | 14 | |
| Plate 3, C5 | M34V, K98E, E122K, T136S, K189E, F205S | A109A | 1.6 | 161 | 17 | 70.6 | 0 | 14 | |
| Plate 7, C4 | T136S, K145M, E156D, D198G | T43T | 1.73 | 169 | 14 | 81.7 | 1.8 | 15 | |
| Plate 3, D5 | K145M | L83L | 1.75 | 228 | 9 | 81 | 0.7 | 11 | |
| Plate 7, H12 | G42S, D198N | | 2.35 | 261 | 3 | 79.8 | 0.6 | 13 | |
| Plate 7, F8 | I111M, I208N | | 1.65 | 175 | 8 | 71.1 | 0.5 | 13 | |
| Plate 7, F9 | E23K, L120M | | 1.55 | 265 | 9 | 83 | 1.3 | 8 | |
| Plate 6, H2 | G104S, D202E | | 2.09 | 237 | 6 | 78.8 | 0.4 | 12 | |
| Plate 5, G12 | K78N, P143T, L212M, Q217L | L47L, L107L | 1.59 | 157 | 4 | 74.4 | 0.6 | 14 | |
| Plate 3, H4 | L47Q, L157M, N194T | D76D | 1.76 | 166 | 12 | 78.8 | 1.1 | 15 | |
| Plate 5, E2 | E62D, H138Q | | 1.5 | 218 | 6 | 78.5 | 0.5 | 10 | |
| Plate 8, F6 | Y52F, A91T, L191Q | | 1.88 | 233 | 6 | 72.3 | 0.2 | 11 | |
| Plate 1, F4 | C130S, I140L | A211A | 0.93 | 69 | 8 | 78.3 | 0.4 | 19 | |
| Plate3, B3 | L191Q | L192L | 0.85 | 188 | 15 | 74.3 | 0.4 | 6 | |
| Plate 3, E6 | N2S, I183F, D198V, Q217R | | 0.63 | 181 | 11 | 68.1 | 0.4 | 6 | |
| Plate 5, D6 | D97V, K101I | | 0.91 | 268 | 40 | 73.4 | 0.8 | 13 | |
| Plate 5, G7 | K189T | H28H, K193K | 0.73 | | | | | | |
| Plate 6, A10 | E22G | | 0.75 | 170 | 6 | 80 | 0.4 | 6 | |
| Plate 6, E1 | P60S | L182L | 0.66 | 114 | 10 | 76.4 | 0.3 | 8 | |
| Plate 7, A10 | L120P, V139A, S169G | K19K | 0.52 | 94 | 9 | 79.9 | 0.8 | 8 | |
| Plate 7, F1 | E102D | | 0.78 | 211 | 21 | 79.7 | 0.5 | 5 | |
| Plate 7, G10 | T64I | A211A | 0.54 | 84 | 12 | 82.4 | 0.2 | 6 | |
| Plate 8, A6 | E44V | L58L, V174V | 0.61 | | | | | | |
| Plate 8, A11 | T89S, R128C | | 0.67 | 69 | 8 | 78.3 | 0.2 | 13 | |

$WT^{syn}$ = WT with synonymous mutations.

**Supplementary Table 6: Activity and thermostability data for AfAdk variants selected in *in vitro* experiments.**

| ID | Variant | Synonymous mutations | Lysate activity (normalized) | $k_{cat}$ ($s^{-1}$) | $k_{cat}$ s.d. | $T_m$ (˚C) | $T_m$ s.d. | Cellular [Adk], mM | Background (vector) mutations? |
|---|---|---|---|---|---|---|---|---|---|
| N/A | WT | | 1 | 134 | 11 | 71.3 | 0.3 | 3 | |
| Plate 1, A5 | R55C, Y109N, A166E | | 1.24 | | | | | | |
| Plate 1, A7 | K23N | | 1.77 | 127 | 7 | 68 | 0.01 | 7 | |
| Plate 1, A12 | I29N | P87P | 1.85 | 152 | 1.3 | 61 | 0.1 | 6 | |
| Plate 1, D1 | S73R | L72L | 1.36 | | | | | | |
| Plate 1, D12 | L45Q, A135V, Q203R | Q48Q | 1.34 | | | | | | |
| Plate 1, G5 | V204I | | 2.61 | 256 | 27 | 71.2 | 0.3 | 5 | |
| Plate 1, G12 | F205L | N164N, A169A | 1.32 | | | | | | |
| Plate 1, H2 | R123L | L120L | 2.11 | 124 | 4 | 67.4 | 0 | 8 | downstream of ori |
| Plate 2, B7 | $WT^{syn}$ | A11A | 1.39 | | | | | | |
| Plate 2, G3 | V204I | A187A | 2.07 | 256 | 27 | 71.2 | 0.3 | 4 | |
| Plate 3, A9 | N79S, P181L, V204I | E94E | 1.7 | 152 | 3 | 68.7 | 0.2 | 5 | |
| Plate 3, B9 | T98I, V204F | | 1.44 | 214 | 19 | 63.6 | 0.3 | 3 | |
| Plate 3, D3 | A206T | A17A, L182L | 1.46 | 133 | 10 | 70.1 | 0.3 | 5 | |
| Plate 1, A10 | N2Y, G134D | L210L, T125T, A145A, G190G | 0.84 | 115 | 10.9 | 70.6 | 0.4 | 4 | |
| Plate 1, C5 | I39F | Q203Q | 0.79 | 120 | 5 | 69.6 | 0.3 | 5 | |
| Plate 1, C9 | $WT^{syn}$ | L45L, L83L | 0.88 | | | | | | YPB1 |
| Plate 1, D6 | A37V, P87A | L3L, E210E | 0.67 | 37 | 1 | 58.3 | 0.1 | 8 | |
| Plate 1, F10 | $WT^{syn}$ | A91A | 0.94 | | | | | | RBS |
| Plate 1, G8 | $WT^{syn}$ | V204V | 0.88 | | | | | | |
| Plate 1, H3 | G190V | A37A | 0.56 | 59 | 6 | 61 | 0.1 | 5 | |
| Plate 2, A3 | D57E, A145V, F205L | | 0.85 | 128 | 9 | 67.3 | 0.2 | 3 | |
| Plate 2, B9 | $WT^{syn}$ | L99L | 0.72 | | | | | | |
| Plate 2, C12 | V68E | | 1.37 | 96 | 7 | 59.7 | 0.3 | 6 | |
| Plate 2, D5 | G42A, L139P, G197D | | 1.13 | | | | | | |
| Plate 2, D7 | WT* | S217S | 0.81 | | | | | | |
| Plate 2, H2 | Q16L, E44K, E97V, N164I | P181P | 0.94 | 85 | 9 | 72.4 | 0.5 | 5 | upstream of bom |
| Plate 3, A11 | G134S | Q48Q | 0.73 | | | | | | YPB1 |
| Plate 3, A12 | P60S, L192V | P87P, D108D | 0.65 | | | | | | upstream of bom |
| Plate 3, B1 | $WT^{syn}$ | R160R, S217S | 1.02 | | | | | | |
| Plate 3, B3 | $WT^{syn}$ | L212L | 0.58 | | | | | | |
| Plate 3, B6 | V68E | | 0.87 | 96 | 7 | 59.7 | 0.3 | 3 | |
| Plate 3, B11 | E132K | | 0.63 | | | | | | rop |
| Plate 3, C5 | N79D | L192L | 0.69 | 104 | 10.4 | 69.6 | 0.3 | 3 | |
| Plate 3, D2 | C130S, Q216* | | 1.31 | | | | | | |
| Plate 3, D4 | R128H, H138Q | L45L, I65I | 0.72 | 109 | 12 | 68.9 | 0.2 | 3 | |
| Plate 3, F2 | L182Q, V204F | I107I, T125T | 0.52 | | | | | | |
| Plate 3, G10 | $WT^{syn}$ | L183L | 1.11 | | | | | | |
| Plate 3, H7 | I195N | T64T, L211L | 0.82 | 86 | 6 | 58.3 | 0.2 | 3 | |

$WT^{syn}$ = WT with synonymous mutations.

### References

1. Billerbeck, S. & Panke, S. A genetic replacement system for selection-based engineering of essential proteins. *Microbial Cell Factories* **11**, 110 (2012).
2. Nguyen, V. *et al.* Evolutionary drivers of thermoadaptation in enzyme catalysis. *Science* **355**, 289–294 (2017).
3. Karst, S. M. *et al.* High-accuracy long-read amplicon sequences using unique molecular identifiers with Nanopore or PacBio sequencing. *Nature Methods* **18**, 165–169 (2021).
4. Steel, H., Habgood, R., Kelly, C. L. & Papachristodoulou, A. In situ characterisation and manipulation of biological systems with Chi.Bio. *PLOS Biology* **18**, e3000794 (2020).
5. Henzler-Wildman, K. A. *et al.* A hierarchy of timescales in protein dynamics is linked to enzyme catalysis. *Nature* **450**, 913–916 (2007).
